# Less is more: Slow-codon windows enhance eGFP mRNA resilience against RNA interference

**DOI:** 10.1101/2024.09.27.615338

**Authors:** Judith A. Müller, Gerlinde Schwake, Anita Reiser, Daniel Woschée, Zahra Alirezaeizanjani’, Joachim O. Rädler, Sophia Rudorf

## Abstract

Extensive efforts have been devoted to enhance the translation efficiency of mRNA delivered to mammalian cells via codon optimization. However, the impact of codon choice on mRNA stability remains underexplored. In this study, we investigated the influence of codon usage on mRNA degradation kinetics in cultured human cell lines using live-cell imaging on single-cell arrays (LISCA). By measuring mRNA lifetimes at the single-cell level for synthetic mRNA constructs, we confirmed that mRNAs containing slowly translated codon windows have shorter lifetimes. Unexpectedly, these mRNAs did not exhibit decreased stability in the presence of siRNA compared to the unmutated sequence, suggesting an interference of different concurrent degradation mechanisms. We employed stochastic simulations to predict ribosome density along the open reading frame, revealing that the ribosome densities correlated with mRNA stability in a cell-type- and codon-position-specific manner. In summary, our results suggest that the effect of codon choice and its influence on mRNA lifetime is context dependent with respect to cell type, codon position, and RNA interference.

## Introduction

Regulating mRNA stability holds significant importance not only for clinical applications^1^ but also within the realm of synthetic biology. Factors that compromise mRNA stability provide potential targets for stability engineering. In this context, the choice of mRNA sequence is particularly important, specifically the mRNA 5’-cap ^2^, nucleotide modifications ^3^, UTR design ^4,5^, poly(A) tail length ^6–8^, RNA interference (RNAi) binding sites ^9^ as well as codon usage ^10,11^. Understanding the underlying mechanisms of these factors is crucial to achieve improved control over the half-life of mRNAs.

Degeneracy of the genetic code enables encoding of the same amino acids via multiple synonymous codons ^12–14^. These synonymous codons are recognized by specific tRNAs and the decoding rate of each one is distinct. The choice of codons determines the translation process and the folding dynamics of the nascent peptide chain ^15^. In nature, genetic codes exhibit codon bias, with diverse organisms displaying varying frequencies of synonymous codons. Optimal codons are those synonymous codons that can be translated more rapidly and accurately ^16,17^. Codon optimality influences translation in several ways, i.e., by changing ribosome speed ^18–23^, translation efficiency ^24–26^, protein folding ^21,27,28^, and translation fidelity ^15,16^. Various computer models have been developed to predict efficient synonymous codon exchange.^29–34^ A comprehensive review can be found in Hanson and Coller (2018) ^15^. Sequence optimization is often performed aiming for enhanced translation ^30,35,36^. Recently, Trösemeier et al. introduced a software for stochastic simulations of mRNA translation ^37^. The software, named OCTOPOS, takes into account many parameters relevant for the translation process to optimize or de-optimize mRNA sequences. To this end, a machine-learning method is applied to combine the simulation results with further mRNA-specific features (such as abundance or length) into a comprehensive model for protein output prediction. In addition to generating optimized mRNA sequences, OCTOPOS calculates steady-state ribosome density profiles, i.e., the ribosome occupancy of individual codons of a sequence.

The effect of ribosome density on translation as well as mRNA stability is not yet fully understood. An inverse correlation between ribosome movement or density along the Open Reading Frame (ORF) and mRNA stability has been observed ^38–41^ and shown to play a critical role in regulating gene expression ^42,43^, and potential ribosome stalling sequences are targeted via no-go decay (NGD) ^44–50^. In contrast, experimental studies indicated that mRNA covered by ribosomes exhibits protection against decay ^51–55^. Deneke *et al*. ^56^ developed a theoretical model that links mRNA degradation and translation based on the assumption that ribosomes protect the mRNA against endonucleolytic degradation processes. Ruijtenberg *et al*. discovered that translating ribosomes play a role in unmasking mRNA, thereby making it accessible for target recognition in RNAi-mediated mRNA cleavage ^57^. Resolving impacts on mRNA stability based on modifications in predicted ribosome occupancy requires precise measurement of translation and degradation kinetics ^58^ However, disentangling mRNA translation and stability in the context of codon usage is experimentally challenging. In previous work it was shown that eGFP reporter translation and lifetime are simultaneously measured using live-cell imaging on single-cell arrays (LISCA) ^59–61^. LISCA monitors the time courses of mRNA-mediated eGFP-fluorescence in hundreds of individual cells on a micro-pattern in parallel. From the dynamics, both mRNA translation and degradation rates are determined based on a kinetic reaction model for mRNA translation including protein maturation as well as protein and mRNA degradation (Fig. 1c) ^60,62^. In previous work Ferizi et al used the LISCA technique to optimize the UTR region of therapeutic mRNA sequences resulting in improved functional mRNA life-time ^4^.

**Figure 1:**
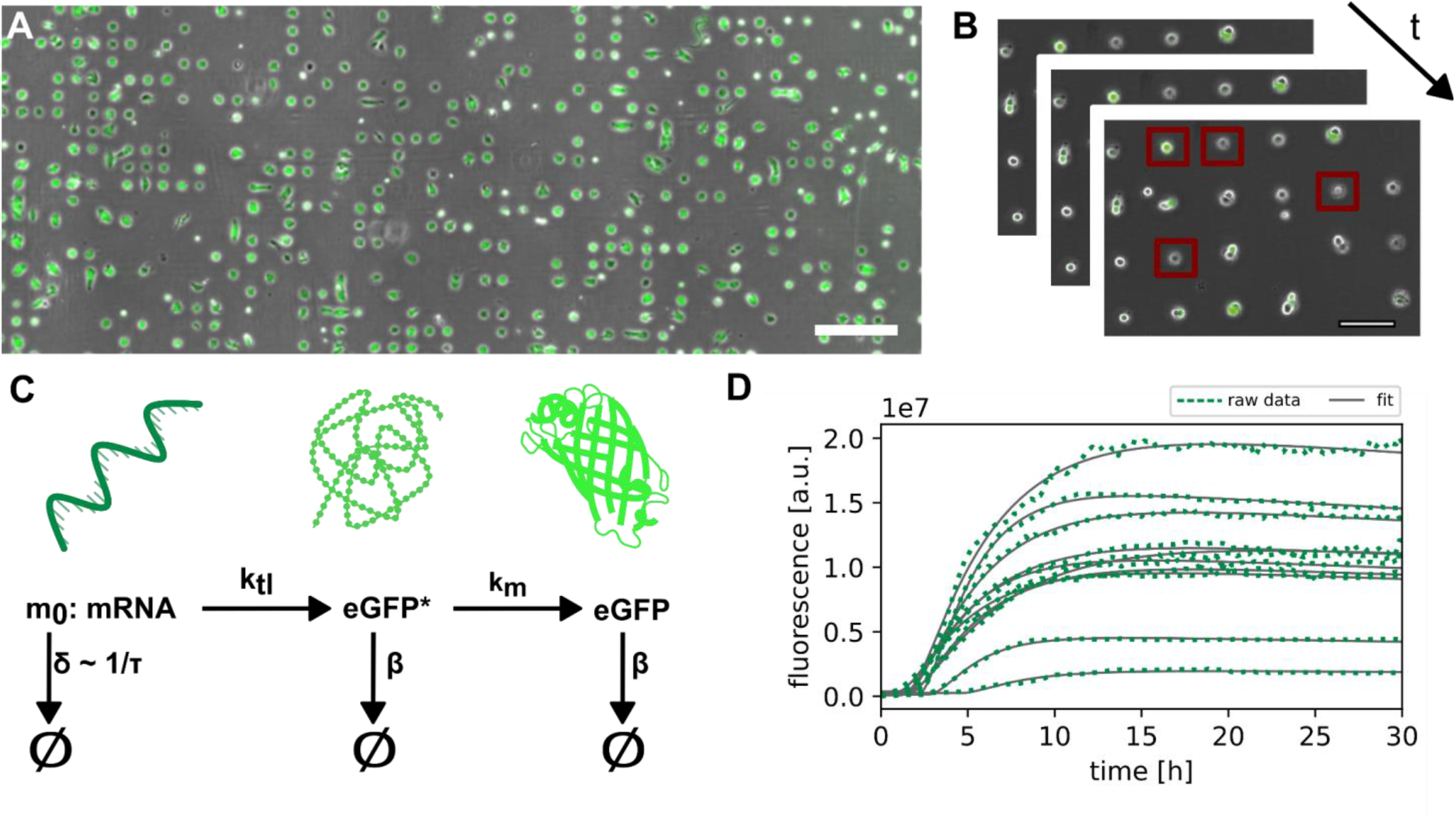
Platform for live-cell imaging on single cell arrays (LISCA): (A) A549 cells on single cell array 24h after transfection. Brightfield and eGFP-fluorescence overlay. Scale bar corresponds to 400 µm. (B) Illustration of the time resolution in single-cell translation experiments with the application of a region of interest (indicated with red squares). (C) Three-stage model for mRNA translation and protein maturation with mRNA stability τ (inverse of mRNA degradation rate δ), translation rate k_tl_, maturation rate km and protein degradation rate ß. (D) Exemplary eGFP fluorescence trajectories of single cells (green) recorded over 30h, fitted to the three-stage translation and maturation model (gray).

Here, we systematically study how slow codon sequences modulate the stability of mRNA constructs with defined siRNA target sites. To achieve this, we designed five constructs of eGFP-mRNA with slowly translated codon windows at different positions within the ORF. We used OCTOPOS to simulate the translation of these constructs and predict their ribosome density profiles.

To assess intracellular stability, we employed LISCA to measure the translation and degradation kinetics of the designed mRNA constructs across hundreds of transfected cells in parallel. Fitting the data to a translational rate model, we calculated degradation and expression rates independently. First, we quantified the decrease in stability of the original eGFP-mRNA construct in the presence of siRNA targeting one of two selected siRNA binding sites. Exact quantification was achieved through co-transfection and normalization with an CayRFP-mRNA control. Next, we examined whether siRNA mediated knockdown is influenced by synonymous codon changes. Correlating the simulated ribosome density profiles with the measured mRNA lifetimes revealed a position-dependent effect of slow-codon windows on mRNA stability. We show that the presence or absence of RNAi changes the effect of codon choice on mRNA lifetime.

## Results

### Live-cell imaging on single-cell arrays (LISCA)

We assess mRNA stability from quantitative analysis of single-cell eGFP gene expression time courses with high temporal resolution. To this end, we acquire hundreds of individual single-cell fluorescence trajectories using the LISCA approach. As in previous work ^60,62^ human lung carcinoma A549 or human liver carcinoma HuH7 cells were seeded in microscopy cell-culture channel slides with micro-patterned surfaces consisting of 10.000 cell-adhesive squares (20×20 µm) allowing for attachment of one cell per spot (**Fig. 1A**). Micro-fabrication was carried out using a refined method (see Methods section and Supplementary Material for details). After an incubation time of 2 hours cells were transfected with reporter mRNAs. Depending on the experimental approach, the mRNA transfection was followed by siRNA transfection. Long-term scanning time-lapse bright field and fluorescence imaging was carried out under physiologic conditions in order to follow eGFP and CayRFP translation dynamics. (**Fig. 1B**). Subsequently, fluorescence time-traces of individual cells were extracted from the image time series using the python based in-house cell tracking software PyAMA ^63,64^ (**Fig. 1D**). Fluorescence time courses were fitted to a kinetic model of protein expression including mRNA translation, mRNA degradation, protein maturation and protein degradation as illustrated in **Fig. 1C&D**. The analysis yields individual values for the kinetic rates per cell for mRNA degradation, δ, and the initial expression rate m_0_*k_tl_, where m_0_ denotes the number of mRNA molecules translated and k_tl_ the translation rate (**Suppl. Fig. S1**). Protein maturation and degradation rate were analyzed in previous work and fixed as described earlier ^60^. This allowed precise fitting of each fluorescence trace as illustrated in **Suppl. Fig. S2**. In the following results, we discuss mRNA stability (or lifetime), τ, as the inverse of the measured mRNA degradation rate δ. In each assay, cells were co-transfected with the mRNA of interest (i.e., one of the eGFP mRNA variants) together with a reference reporter mRNA (Cayenne red fluorescence protein, CayRFP). Co-transfection analysis of eGFP and CayRFP fluorescence results in a distinct eGFP and CayRFP fluorescence trajectory for every single cell. This then allows us to normalize the eGFP mRNA parameters with the respective CayRFP mRNA parameters of the same cell as explained in **Suppl. Fig. S3**. Thereby, we address intrinsic cell-to-cell variance in expression levels caused by varying numbers of lipoplex particles delivered per cell as well as varying cell-cycle or metabolic state of each cell. All kinetic rates are reported in terms of fold-change of eGFP mRNA kinetic rates normalized to the CayRFP mRNA reference value (**Suppl. Fig.S3 and S4)**. mRNAs were premixed before lipoplex formation in order to secure equal amounts of mRNA molecules per lipoplex ^65^. As previously reported, the two-color single-cell referencing approach enhances the accuracy of measurement of mRNA degradation and expression rates yielding an improved signal-to-noise ratio up to a factor 5 compared to population measurements ^66^. As control, single cell correlation analysis was performed. No correlations were found in eGFP and CayRFP mRNA degradation rates, while some correlations were observed in initial eGFP and CayRFP mRNA translation rates m_0_*k_TL_ (**Suppl. Fig S5)**. Latter are the result of eGFP/RFP mRNA codelivery and the dominating influence of lipoplex number fluctuations that determine the amount of initial number m_0_ of mRNA released.

### Quantification of RNAi mediated mRNA decay

Specific and targeted triggering of mRNA degradation using siRNA allows for quantification of RNAi mediated mRNA decay as depicted schematically in **Fig. 2A**. We performed LISCA on cells transfected with siRNA in addition to eGFP- and RFP-mRNA. The siRNA was targeted against the ORF of eGFP mRNA with binding sites at nucleotide position 122 (siRNA 1) and 433 (siRNA 2), respectively, see supplementary information for details. A non-binding siRNA (siCtrl) was used as reference control. To confirm the specificity of eGFP-mRNA targeting over RFP mRNA, we assessed their stabilities in the presence and absence of siRNA. **Fig. 2B-D** shows siRNA induced knockdown in terms of eGFP-mRNA life-time reduction. The lifetime of eGFP mRNA is reduced in the presence of both siRNAs compared to siRNA control (**Fig. 2B&C**), with a median fold change of normalized eGFP mRNA stability of 0.11±0.07 for siRNA 1 and 0.08±0.05 for siRNA 2 in A 549 cells. In HuH7 cells, the difference between both siRNAs was found to be higher: A fold-change in stability of 0.09±0.02 was found for siRNA 1 whereas transfection of siRNA 2 led to a fold-change of 0.25±0.07. In contrast to the eGFP mRNA stability, the CayRFP reference construct’s stability remains unaffected with siRNA 1 or siRNA 2 (see **Fig. 2B&D**). Notably, the normalized initial expression rates m_0_*k_tl_ are not influenced by the presence of eGFP mRNA specific siRNA (see **Fig. 2C&E**) and no correlation exists between expression rates and mRNA stability at the single-cell level (**Fig. 2C&E, Suppl. Fig. S1**), or between eGFP- and RFP-mRNA (**Suppl. Fig. S5**). These results demonstrate the capability of LISCA to quantitatively measure mRNA stability independently of translational speed and that RNAi can serve as a suitable tool to study mRNA stability in LISCA experiments.

**Figure 2:**
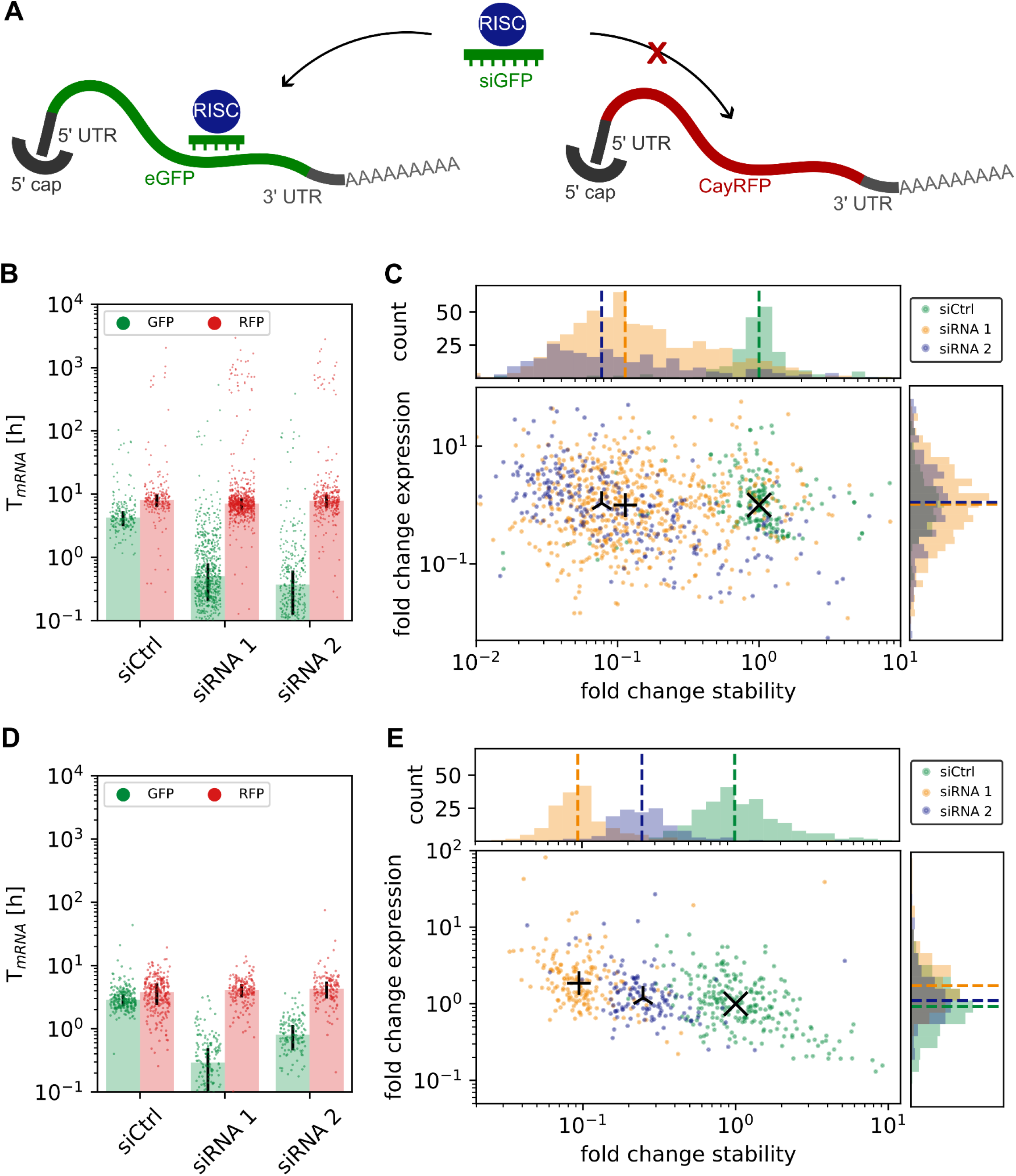
Determination of mRNA stability at the single-cell level: (A) RNAi mediated degradation by siRNA targeted specifically against the ORF of eGFP but not CayRFP mRNA. (B, D) Single-cell mRNA half-life (dots) alongside median half-life (bars) with MAD (error bars), demonstrating the decrease in eGFP mRNA stability in the presence of siRNA 1 or 2 but not siCtrl (non-binding control), with no impact on the stability of co-transfected CayRFP mRNA. (C, E) Scatter plots and corresponding histograms show decrease of median mRNA stability but constant initial mRNA expression rate compared to the median of experiments with siCtrl. Individual data were normalized to respective CayRFP mRNA values for each cell. (B) and (C) show data measured in A459 cells, (D) and (E) respective data for HuH7 cells.

### Generation of predicted slow-codon windows

Next, we study the influence of non-optimal codons in the ORF on mRNA half-life in the context of RNAi. Prior to target mRNA cleavage, a siRNA/RISC complex needs to bind to the target mRNA and find the cleavage site. Ribosomes and the translational machinery are known to interfere with this process ^57,67–69^. Here, we aimed to modulate ribosome density in a site-specific manner and investigate the effect on mRNA stability under siRNA attack This was achieved by synonymous codon exchange. Depending on their optimality or de-optimality, codons are translated at different rates, leading to modulations of ribosomal speed along the ORF (**Fig. 3A**) ^11,70,71^. A stretch of several consecutive non-optimal, slow codons was described to cause a local increase in ribosome density, especially if those codons are preceded by codons with higher optimality^72^. Due to the stochastic nature of translation elongation, ribosome jams or collisions might occur ^38,45,47,72^. Thus, one way to control translation and thereby ribosome speed on the ORF is to replace individual codons with synonymous but more slowly translated non-optimal codons. We simulated 230 synonymous variants of eGFP mRNA, each containing a window of 10 adjacent non-optimal but synonymous codons at a different position. Using our software OCTOPOS ^37,73^, we simulated ribosome movement on the ORF of these 230 variants and identified 5 sequences with high similarity in terms of overall ribosome flux (**Suppl. Fig. S6**). Predicted ribosome densities along the ORFs of these 5 constructs compared to unmutated eGFP mRNA are shown in **Fig. 3B**. The simulations show an increased ribosome density for all constructs with the peak occupancy in proximity of the non-optimal codon window. Changing the translation initiation rate in the simulations within physiological limits alters the shape of the resulting ribosome density profiles but not the overall effect (**Suppl. Fig. S7**).

**Figure 3:**
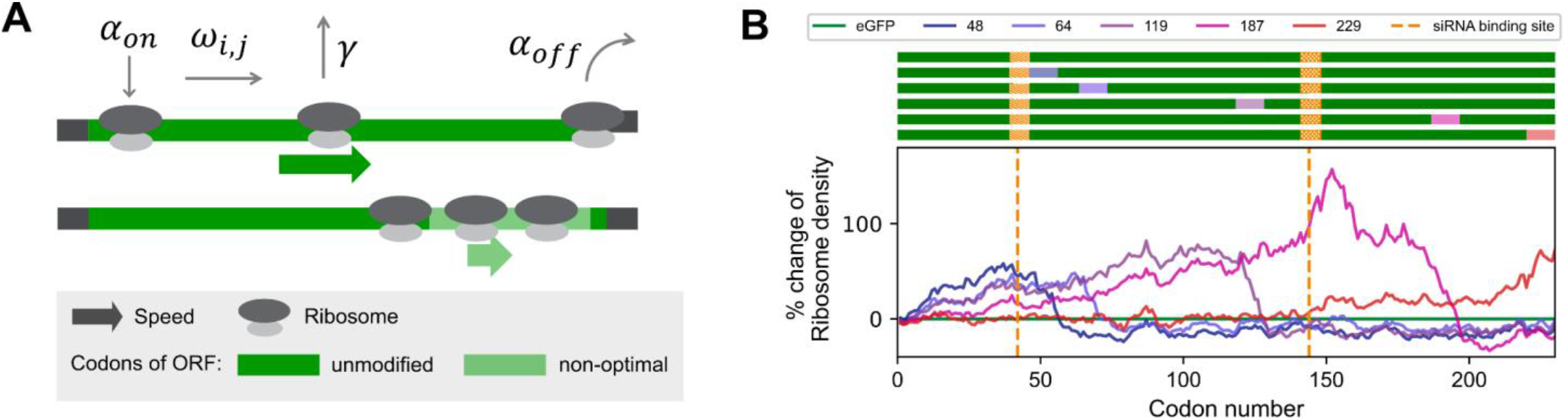
Computational predictions of ribosome density modulations on the ORF of eGFP mRNA: (a) Scheme of the modeling concept with binding rate α_on_, elongation rates ω_i,j_ of codons i in mRNA j, premature drop-off rate γ, and translation termination rate α_off_. Replacement of stretches of original codons (dark green) by their synonymous but non-optimal alternatives (light green) is assumed to cause a slow-down of translation at these slow-codon windows, which can lead to a local increase in ribosome density. (b) Top: ORF of eGFP mRNA variants (dark green) with slow-codon windows (colored boxes at indicated positions) and siRNA binding sites (orange boxes). Bottom: Predicted differences of ribosome density profiles along the ORF of unmutated eGFP (green) and variants with slow-codon windows at indicated positions (purple to red). Numbers indicate the position of the first inserted slow-codon.

### mRNAs with non-optimal codon windows show decreased stability

It was described previously that - in absence of any siRNA - locally increased ribosome densities can potentially trigger cellular quality control mechanisms leading to mRNA degradation ^11,22,72^. To test this within our single-cell experimental approach, we transfected A549 and HuH7 cells, respectively, on a single-cell array with mRNA encoding either eGFP or one of the five synonymous variants featuring slow-codon windows, as outlined in the previous section. CayRFP as internal reference was co-transfected. By fitting the three-stage maturation model to the experimentally determined single-cell fluorescence trajectories, we assessed the stability of the variant mRNAs. We standardized these results using the co-transfected reference CayRFP mRNA and normalized them to the corrected stability of the unmutated eGFP mRNA, as explained earlier. All constructs exhibited either similar or decreased stabilities compared to unmutated eGFP mRNA (**Fig. 4A&C, Suppl. Fig. S8** for single-cell data). Mann-Whitney U tests revealed significant changes for all constructs except for the mRNA with a slow-codon window positioned near the end of the ORF (codon positions 229 to 239) in the A549 cell line (**Fig. 4A**). A similar, although not identical trend, was observed in the HuH7 cell line. Here, we also observed mostly impaired stabilities (see **Fig. 4C**). The most substantial impact on mRNA stability, reducing it to 66% ± 7% compared to unmutated eGFP mRNA stability, was observed for the mRNA with a slow-codon window from position 64 to 74 in the A549 cells. The strongest reduction in HuH7 cells was observed for the slow-codon window starting from position 119, with a relative stability of 71% ± 6%. It’s noteworthy that the initial expression rates m_0_*k_TL_ also varied compared to unmutated eGFP mRNA, but with no discernible trend toward overall increased or decreased expression in neither of the tested cell lines, as illustrated in **Suppl. Fig. S9A&C** (**Suppl. Fig. S8** for single-cell data).

**Figure 4:**
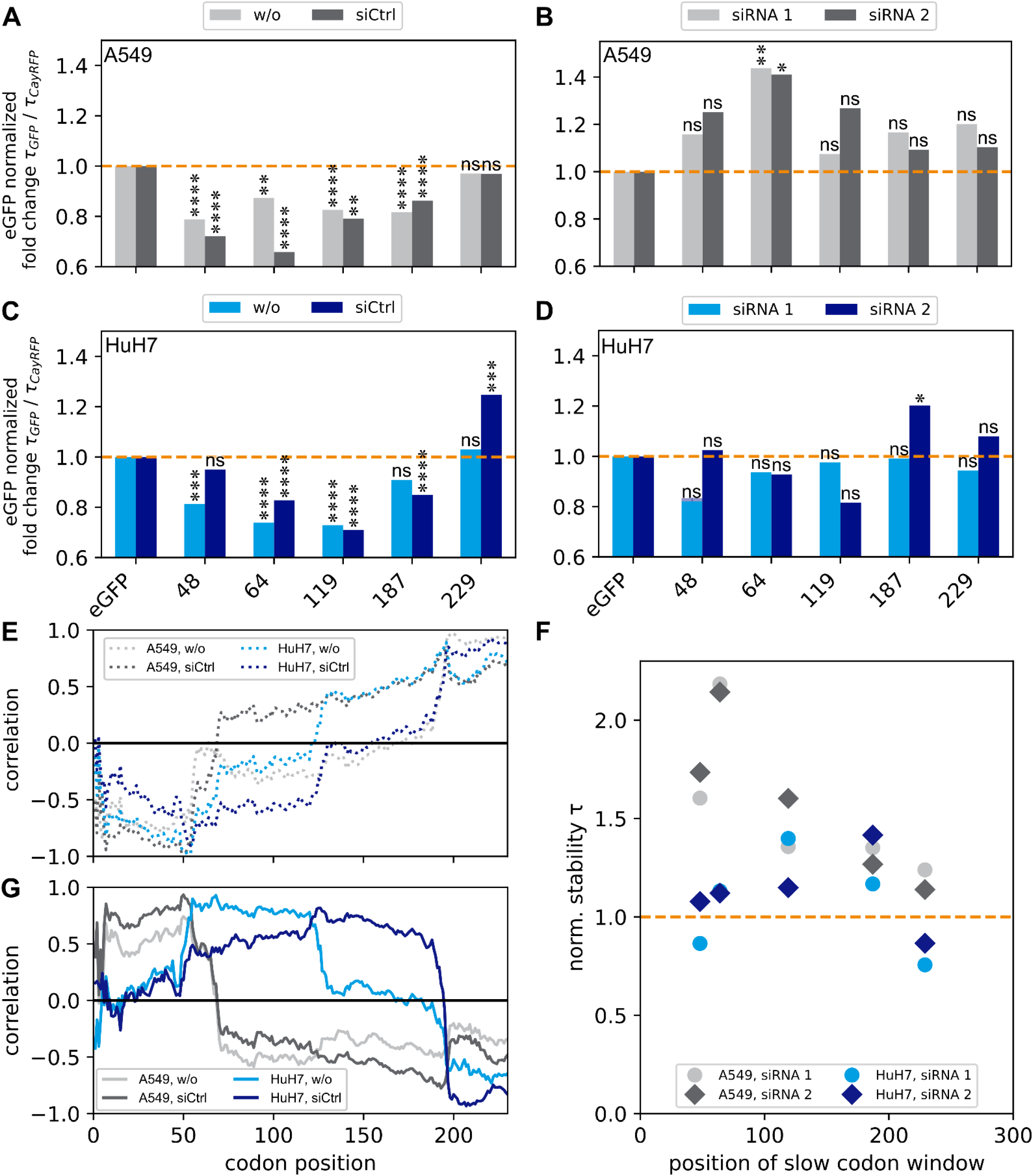
Destabilization vs. Stabilization – Effect of non-optimal codons on mRNA stability: Stability of eGFP mRNA was determined by LISCA and normalized with equivalent values from reference CayRFP mRNA for each cell. (A) Almost all constructs with non-optimal codon windows have a decreased median stability in control experiments without siRNA (w/o) or with control siRNA (siCtrl) in comparison to unmodified eGFP mRNA in A549 cells. (B) When siRNA 1 or 2 is added, the median stability compared to eGFP is rescued compared to the control experiment in A. Significance levels were assessed using the Mann-Whitney U-test. Detailed single-cell data are provided in the Supplementary Information. (C) and (D) equivalent data acquired in HuH7 cells. (E) Correlation of measured mRNA stability with simulated ribosome density is largely independent of cell type. (F) NormalizedmRNA stabilities under RNA interference for siRNA 1 (dots) and 2 (diamonds) for A549 (gray) and HuH7 (blue) cells. Normalization with respect to mRNA stabilities obtained under addition of siCtrl. (H) Correlation of normalized stability and predicted ribosome density on the five constructs for each codon position along the ORF. See also Suppl. Fig. S10 for more details.

### siRNA mediated degradation mitigates the stability disadvantage of mRNAs with non-optimal codon windows

To test the effect of slow-codon windows on the effectiveness of RNAi mediated degradation, we transfected A549 and HuH7 cells with CayRFP and either unmutated eGFP mRNA or one of the slow-codon window variants, followed by transfection with either negative control siRNA (siCtrl), or targeting siRNA variant 1 or 2. We determined the stability of the transfected mRNAs for each cell by LISCA, where we normalized both to the stability of the transfection control CayRFP mRNA and of the unmutated eGFP mRNA experiment, see **Suppl. Fig. S8** for single-cell data. As expected, siCtrl does not have a systematic effect on mRNA stability and we obtained similar results as in the absence of any siRNA, i.e., constructs with slow-codon windows are less or equally stable than unmutated eGFP mRNA (**Fig. 4A&C**). Additionally, we computed the correlation coefficient between the measured stabilities and the simulated ribosome densities for each codon position within the ORF. This analysis revealed consistent correlation profiles for both cell lines when no siRNA or siCtrl was transfected. Specifically, we observed a strong negative correlation between simulated ribosome density and mRNA stability at the 5’ end of the ORF, which transitions to a positive correlation towards the 3’ end (**Fig. 4E**). These results suggest that higher ribosome occupancy at the beginning of the ORF negatively impacts mRNA stability, whereas higher ribosome density at the end of the ORF is beneficial. Introducing siRNAs targeting the eGFP mRNA constructs leads to an overall reduction of stability for all tested constructs (**Suppl. Fig. S10**). When comparing the fold-change stabilities of the mutated constructs with unmutated eGFP mRNA, we observe that the constructs with slow-codon windows are now as stable as or more stable. Importantly, unlike the scenario without functional siRNA, where we observed a significant decrease in stability, we now mostly observe insignificant changes (**Fig. 4A-D**). This indicates that the stability of mutated and unmutated constructs is at least equal under RNA interference conditions, and for individual cases we even find a significant stabilization. This means that RNA interference diminishes the stability of mRNAs containing slow-codon windows by a smaller factor compared to unmutated eGFP mRNA.

### mRNAs with non-optimal codon windows are less affected by siRNA mediated degradation

To better assess the relative stabilization of mRNAs under RNA interference by slow-codon windows, we further normalize mRNA lifetimes measured in presence of siRNA 1 or 2 by the respective median values obtained in the presence of non-functional siCtrl. By doing so, it is possible to examine the relative influence of RNA interference on mRNA stability in isolation regardless of other effects that might influence the stability such as quality control pathways or transfection associated effects. We found a relative stabilization of up to 2.2±0.4 fold in A549 cells and up to 1.4±0.2 fold in HuH7 cells, see **Fig. 4F**. However, the normalized mRNA lifetimes correlate only partially with simulated ribosome densities at the specific siRNA binding sites, see **Suppl. Fig. S11**. Notably, we also observed increased relative stabilities of mRNAs with non-optimal codon windows positioned upstream of the binding site of siRNA 2, which were predicted to not have increased ribosome densities at the siRNA binding site in our simulation. Therefore, we again determined the coefficient of correlation of the strength of stabilization and ribosome density for each codon position along the ORF, see **Fig. 4G** and Suppl. **Fig. S12**. Here, we found that higher ribosome occupancy provides greater protection against siRNA-mediated degradation within the first 70 codons in A549 cells, and similarly for the major part of the ORF in HuH7 cells. Conversely, towards the 3’ end, the simulated ribosome densities show an anti-correlation with experimentally determined mRNA stability.

In summary, our findings indicate that under conditions where mRNA is subjected to siRNA/RISC interference, non-optimal codon windows exhibit a protective effect, despite causing a reduction in stability in the absence of functional siRNA. For the two examined siRNAs, this protective effect is largely independent of the targeted cleavage site but seems to depend on the position of the non-optimal codon window and to be positively correlated with the ribosome density in the first part of the ORF, with some cell-type specific variations. The combination of LISCA and stochastic simulation makes the signature of position-dependent correlations accessible, providing interesting new approaches to studying the molecular details of RNA interference.

## Discussion

In this study, our objective was to elucidate the complex relationship between codon optimality and mRNA stability. We utilized live-cell imaging on single-cell arrays (LISCA) to evaluate mRNA stability and expression rates of various exogenous eGFP-mRNA reporters versus CayRFP-mRNA reference systems. We observed a consistent decrease in mRNA lifetime for five mutated eGFP mRNA constructs containing slow-codon windows at specific sites compared to the original eGFP reporter. In the presence of targeting siRNAs, however, the knockdown effect was seemingly attenuated by the introduction of slow codons, resulting in overall stabilities comparable to unmutated eGFP mRNA.

A possible explanation of these findings is that suboptimal codons stabilize mRNA integrity against RNA interference, mediated via modulation of the ribosome density profile. Alternatively, our results may indicate the interference of different mRNA degradation mechanisms when both slow-codon windows and siRNA are present. The regulation of both endogenous and exogenous mRNA stability is a complex process, influenced in part by the translation process itself ^11,74^. As mRNA translation is a stochastic process, ribosome collisions can occur and trigger rescue mechanisms ^38,47,48,50^. As previously noted in various studies, synonymous codon exchange within the ORF comes with a well-described adverse effect of codon suboptimality on mRNA lifetime ^11,22,41,43,49^. In contrast, for *S. cerevisiae* a protective effect was reported for slow-codon windows that are located close to the 5’ end of the ORF and potentially decrease the density of ribosomes further downstream ^75^.

We confirmed the destabilizing effect of slow-codon windows on eGFP mRNA in two different human cell lines and found it to be position-dependent. Based on simulated ribosome density profiles, we assume that the translational machinery is slowed down in a position dependent manner. The introduced changes in ribosomal density may then mediate both, the triggering of rescue mechanisms and a potential protection against siRNA-RISC attack. Moreover, we computed the correlation of predicted ribosomal densities and measured mRNA stability for all codon positions in the ORF, where the first part of the ORF appeared as a particularly sensitive region in both tested cell lines. A high predicted ribosome density within this region is positively correlated with a low mRNA stability in general, but has a seemingly protective effect against siRNA-mediated degradation. This observation is independent of the position of the siRNA binding site. Differences between the tested cell lines are expected, e. g. due to differences in gene expression and metabolic characteristics of both cell types. However, it remains an open question which mechanisms lead to the observed correlation profiles of ribosome density and mRNA stability under RNAi. RNAi mediated degradation is initiated by binding of the siRNA seed region ^67,76^. Therefore, we mapped potential seed binding regions on the ORF for siRNA 1 and 2 to investigate if this explains the observed position dependence, and we indeed found an increased frequency of siRNA seed binding regions towards the 5’-end (**Suppl. Fig. S12**). Previous studies showed a close interplay between ribosome movement and RISC binding (43, 44). In principle, mRNA secondary structure could impact the effects described here ^26^. Through the literature, secondary structure is discussed to affect RISC target accessibility as for example described by Ruijtenberg et al. ^57^, Brown et al. ^77^ or Ameres et al.^68^. A theoretical structure prediction using RNAfold ^78^ for the five eGFP constructs did not provide any hints towards this direction (**Suppl. Fig. S13**), although we cannot ultimately rule out that changes in mRNA stability under RNAi are caused by altered secondary structures. However, focusing on just the secondary structure of an mRNA neglects the potential role of ribosomes. Ruijtenberg *et al*. described how translating ribosomes de-mask RISC binding sites on the mRNA.^57^ This mechanism can explain the reduced stability for the unmutated eGFP sequence but not the beneficial effect of the slow-codon windows. In summary, the ability to observe protein expression trajectories of hundreds of single cells in parallel by LISCA together with simulations of mRNA translation by our software OCTOPOS allows us to demonstrate that translation dynamics can be manipulated via synonymous codon choice to actively control and modulate mRNA stability. We uncovered that siRNA mediated decay can mitigate the stability disadvantage of mRNAs containing non-optimal codon windows. This approach could be exploited in future applications of exogenous mRNAs especially in potential co-delivery systems.

## Acknowledgments

This work was funded by the Bayerische Forschungsstiftung (Project ID AZ-1350-18) in a collaborative grant with ethris GmbH (Planegg, Germany) and ibidi GmbH (Gräfelfing, Germany).

## Author contributions

**Judith A. Müller** conceived the study, designed, performed and analyzed experiments and wrote the manuscript. **Gerlinde Schwake** performed experiments and developed the platform. **Anita Reiser** performed the initial experiments and contributed to the development of the platform. **Daniel Woschée** wrote the in-house software for data analysis. **Zahra Alirezaeizanjani and Sophia Rudorf** performed ribosome density simulations. **Joachim O. Rädler and Sophia Rudorf** conceived the study, supervised the project and wrote the manuscript.

## Materials and Methods

### Cell culture

A549 cells (DSMZ, AC107) were cultured in RPMI medium (Roswell Park Memorial Institute 1640, Gibco™, Thermofisher Scientific, # 11875093) supplemented with 10% (v/v) FBS (Fetal bovine serum, # 10270106) at 37°C and 5% CO_2_. For HuH7 cell culture, additionally, 5mM HEPES (Gibco™, Thermofisher Scientific, #15630080) and 1mM Na-Pyruvate (Gibco™, Thermofisher Scientific, #11360070) were added. For live-cell imaging, A549 cells were cleaved with T/E (Trypsin/EDTA, Gibco 15400-054), HuH7 cells with Acutase (invitrogen, 00-4555-56), and seeded in growth medium at a cell density of 1*10^6^ cells/mL (A549) or 5*10^5^ cells/mL (HuH7) with 25 µL per channel of the 6-channel µ-slide (ibidi, #80600). After 60 min, cells were washed with OptiMEM (Reduced Serum Minimal Essential Medium, Gibco™, Thermofisher Scientific, #31985062) which was also the medium for any transfection experiment. Cell lines were tested for mycoplasma contamination before experiments and were found to be negative.

### Single cell array fabrication

For the preparation of the single-cell pattern, photo induced CuSO_4_ Click Reaction was carried out to selectively bind the cell-adhesive cyclo-Arg-Gly-Asp (RGD) to a cell repellent PVA surface. Therefore, each channel of a bioinert, PVA coated, µ-slide (ibidi, #80600) was filled with 33 µL of a 5 mM Diazirin (Enamine), 10% (v/v) DMSO (Thermofisher Scientific, # D12345) solution. The slide is illuminated with an in-house UV-illumination lamp (Rapp, 365 nm). To enable selective illumination, a silica-wafer based photomask with 20 µm x 20 µm squares and 85 µm spacer was used. Following a washing step, the click reaction solution was applied (10 mM BTTA (JenaBioscience, CLK-067-25), 2 mM CuSO_4_ (JenaBioscience, CLK-MI005), 0.1 mM cyclo-RGD-azide (Lumiprobe, A1330), 100 mM vitamin C (JenaBioscience, CLK-MI005) in sodium-phosphate buffer (JenaBioscience, # CLK-073)) for 1 h at room temperature. Click mix was removed and the slide was washed with phosphate buffered saline (PBS, Biochrom GmbH #L182) several times. See also **Suppl. Fig. S14**.

### mRNA and siRNA

eGFP constructs were kindly provided by ethris GmbH, sequences of the ORFs are provided in the supplementary information. CayRFP mRNA was produced as described previously in ^60^. mRNA was produced without modified nucleotides. siRNAs were purchased from Dharmacon (P-002048-01-20). Sequences are available in the Supplementary Information.

### Transfection assay

To investigate translation kinetics, cells were transfected with equal amounts of CayRFP-mRNA and the respective eGFP-mRNA. Therefore, mRNAs were mixed with Lipofectamine2000 (ThermoFisher, #11668019) according to the manufacturer’s protocol. 2 ng/µL mRNA concentration transfection mixed was applied and cells were incubated under physiologic conditions for 45 min. Afterwards, cells were washed in OptiMEM and 12.5 pmol of the respective siRNA was applied for 30 min. Then, cells were washed in L15 medium (Gibco™, #21083027), supplemented with 10% (v/v) FCS.

### Fluorescence microscopy

Time resolved fluorescence images were recorded with a Nikon TI Eclipse microscope. To allow image acquisition over 30 h, cells were incubated with the Oku-lab incubation system (cage incubator with active humidity and temperature control) under physiologic conditions. Channels were scanned in a 10 min time interval with 10x magnification (Nikon objective, MRH00101). BF illumination was carried out with a 100W warm white LED (MHLED100W), Fluorescence with a LED light source (lumencor, SOLA-SE II). eGFP fluorescence was captured with the eGFP Filterset (Chroma, F46-002), CayRFP Fluorescence with the DsRed ET Filterset (Chroma, F46-005). Images were captured with a CMOS camera (PCO, pco.edge4.2). Acquisition control was performed with the NIS-Elements Advanced Research software (Nikon).

### Image processing and analysis

Images obtained from the NIS software were converted into time-resolved “.tif” stacks. With the in-house analysis software PyAMA (python based automated microscopy analysis)^63^, cell segmentation, tracking and background correction based on Schwarzfischer^64^ was performed. To improve the signal to noise ratio, integrated fluorescence over a square region of interest with a side length of 350% of the square side length was analyzed.

### Calibration for absolute protein numbers

Conversion of fluorescence intensity into protein numbers was carried out with a PDMS calibration chip with known channel dimensions and protein solutions in different concentrations as described previously^60^.

### Data fitting

A non-linear least square fit of data to a three-stage maturation model (see **Fig. 1c**) for mRNA translation was performed. This system is described with the following system of differential equations:

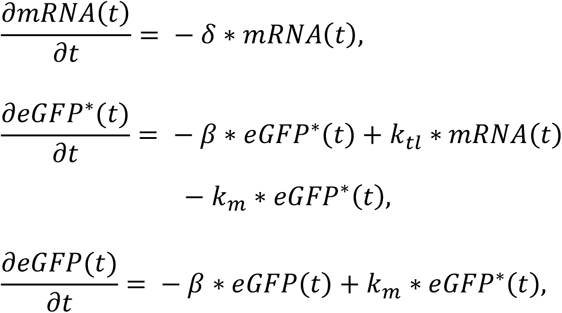

where *mRNA*(*t*) is the concentration of mRNA, *eGFP*\*(*t*) the concentration of nascent eGFP, *eGFP*(*t*) the concentration of maturated eGFP, *δ* the degradation rate of mRNA, *β* the degradation rate of nascent or maturated eGFP, *k*_*TL*_ the translation rate of eGFP encoding mRNA, and *k*_*m*_ the maturation rate of eGFP.

Independent measurement of k_m_ and ß according to Krzyszton *et al*. (2019) ^60^ via induction of translational stop with cycloheximide allowed to reduce the set of free parameters and thus the determination of mRNA degradation and expression rate by model fitting.

### Data normalization and Error calculation

If not stated differently, each experiment was repeated at least three times independently.

All single cell data presented here are normalized to the co-transfected reference mRNA. Therefore, for each single cell *n*, eGFP stability *Г*_*eGFP*_ was normalized to respective CayRFP stability *Г*_*CayRFP*_ of the same cell. 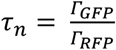 and accordingly for the expression rates:

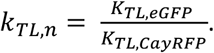

To compare stabilities or expression rates of mutated constructs with corresponding values of unmutated eGFP, data for all separate constructs *i* in each experimental condition *j* (w/o, siCtrl, siRNA 1, siRNA 2) were normalized to the median eGFP mRNA stability in the same experimental condition: 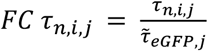. Those values are presented in the graphs unless stated otherwise. For simplicity, only the formula for stability is shown here. Accordingly, expression rates were normalized where applicable.

Figure 4d aimed to compare the RNAi effects only. Therefore, for every single cell n in siRNA j experiments (*j* = 1, 2), fold change *FC τ*_*n,i,j*_ of mRNA stability in presence of siRNA j = 1 and j = 2 was normalized to the corresponding median fold change 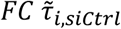 of mRNA stability in presence of siCtrl:

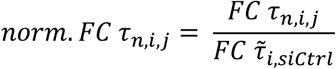

P-values correspond to results from Mann-Whitney-U tests and are indicated in the plots as assigned in the following: (****) for p < 10^−4^, (***) for p < 10^−3^, (**) for p < 10^−2^, (*) for p < 5*10^−2^ and (ns) for p > 5*10^−2^.

