## Supplementary Information for "Less is more: Slow-codon windows enhance eGFP mRNA resilience against RNA interference"

[1] Faculty of Physics, Ludwig-Maximilians-Universitaet, Munich, Bavaria, 80539, Germany
[2] Independent researcher
[3] Leibniz University Hannover, Institute of Cell Biology and Biophysics, 30419 Hannover, Germany


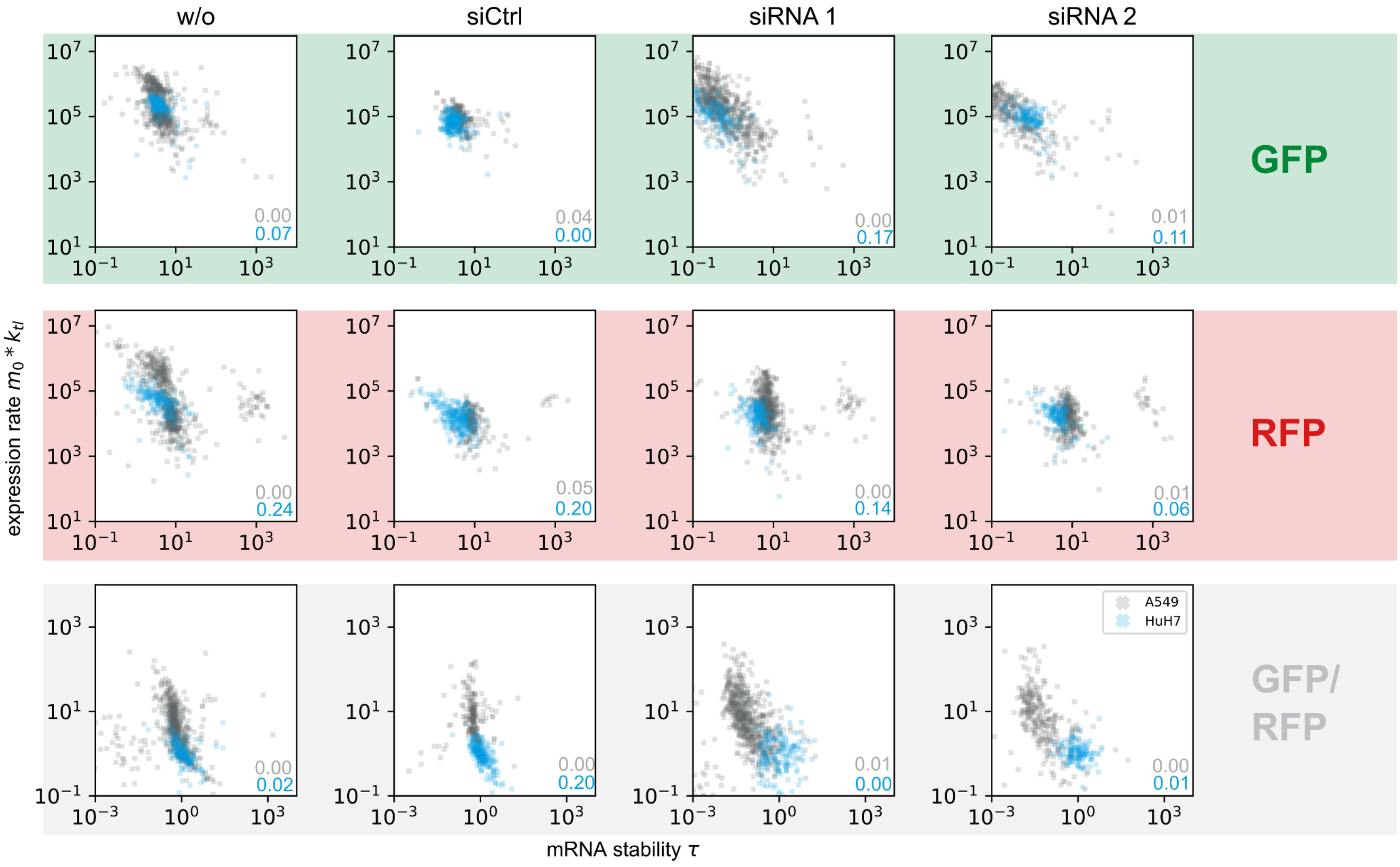


***Figure S1: Initial translation rate $m_0*k_{tl}$ and stability $\tau$ for eGFP and CayRFP mRNA:*** *w/o: without siRNA addition, siCtrl: negative control siRNA, siRNA 1 and 2: addition of eGFP targeting mRNA. Top row (green) shows values for eGFP fluorescence, middle row (red) values for CayRFP fluorescence and bottom row (gray) shows single-cell eGFP data normalized to internal CayRFP reference. Gray crosses correspond to A549 cells, blue crosses to HuH7 cells. $R^2$ values are indicated with the respective color.*

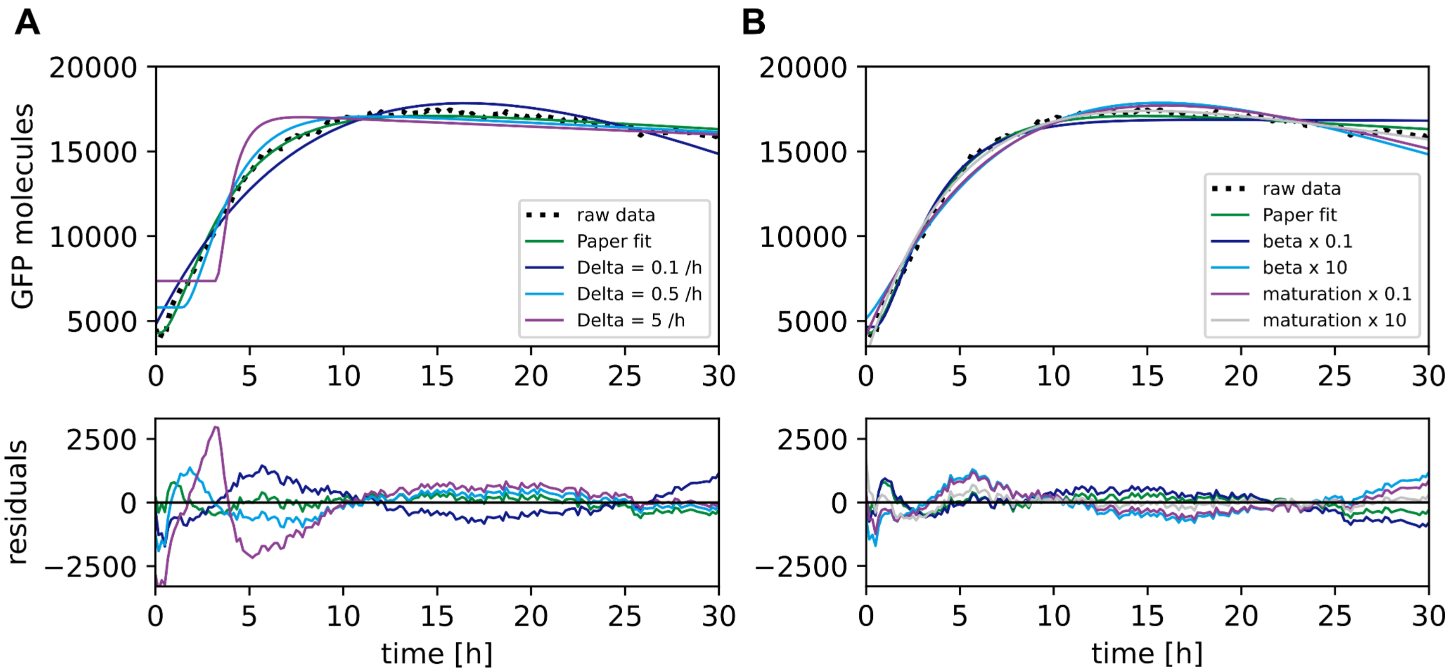


*Figure S2: Fitting of fluorescence traces:* *(A) Exemplary fluorescence trace fitted to the three-stage maturation model described in main* ***Fig. 1C****. Black dots represent raw data. Green line fit with fixed values for protein maturation according to Krzyszton et al.* [1]*. To illustrate the effect of different stability values on the fit, different values for the mRNA stability δ were inserted to the fit (see blue, cyan and purple fits). (B) Corresponding approach but for protein degradation β and protein maturation shows that the shape of the trajectory is less sensitive to the protein specific parameters than to the mRNA specific parameter.*

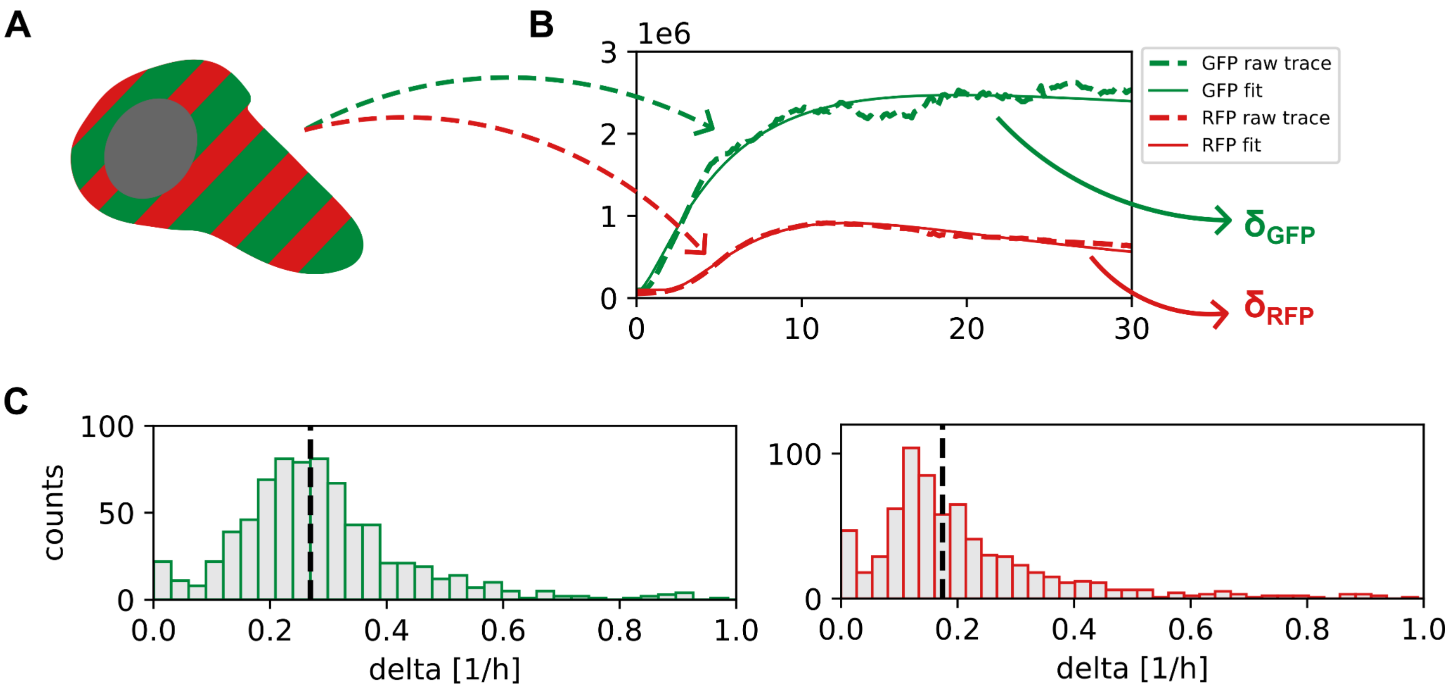


*Figure S3: Single cell normalization:* *(A) Co-transfection of GFP and CayRFP results in cells expressing both proteins. (B) Time-lapse microscopy yields a distinct fluorescence trajectory for each protein per cell. Fitting then results in a distinct values for GFP mRNA degradation ($\delta_{GFP}$) and CayRFP mRNA degradation ($\delta_{RFP}$) (C) Every experimental set resulted in a distribution of fit parameters for all single cells for both mRNA types and allowed cell-wise normalization of all parameters.*

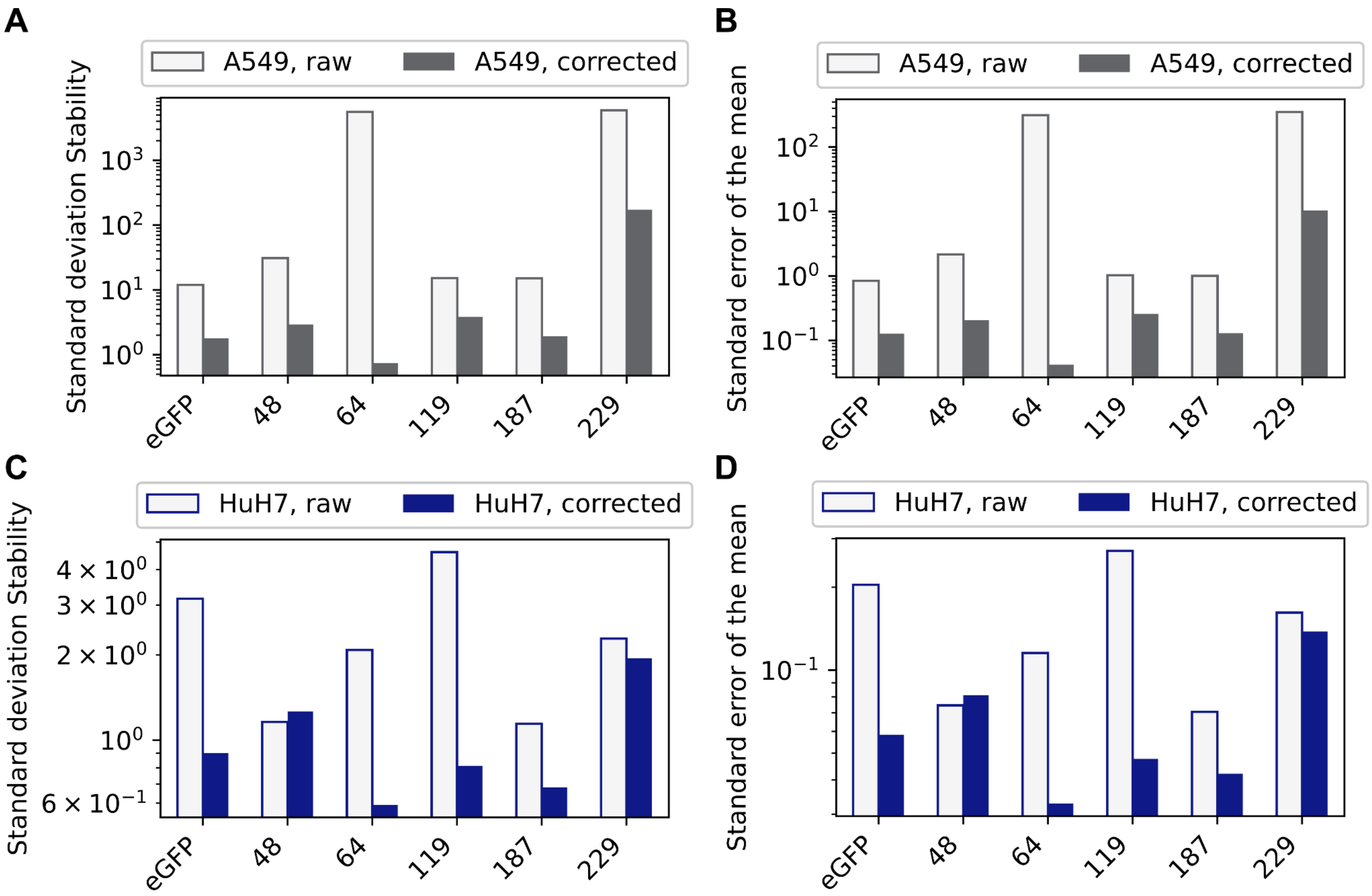


***Figure S4: Single-cell normalization of stability to co-transfected Cay-RFP mRNA leads to reduced standard deviations and standard errors of the mean:*** *For each single cell, the calculated stability of eGFP mRNA was normalized to the stability of CayRFP mRNA. Cells were transfected with both mRNAs with 50 µg per channel, resulting in a total concentration of 2 ng/µL mRNA. Shown here are the standard deviations of stability for eGFP mRNA alone (white bars) and normalized to co-transfected Cay-RFP mRNA (filled bars) for all single cells. (A) Standard deviation in A549 cells, (B) standard error of the mean in A549 cells, (C) standard deviation in HuH7 cells, (D) standard error of the mean in HuH7 cells.*

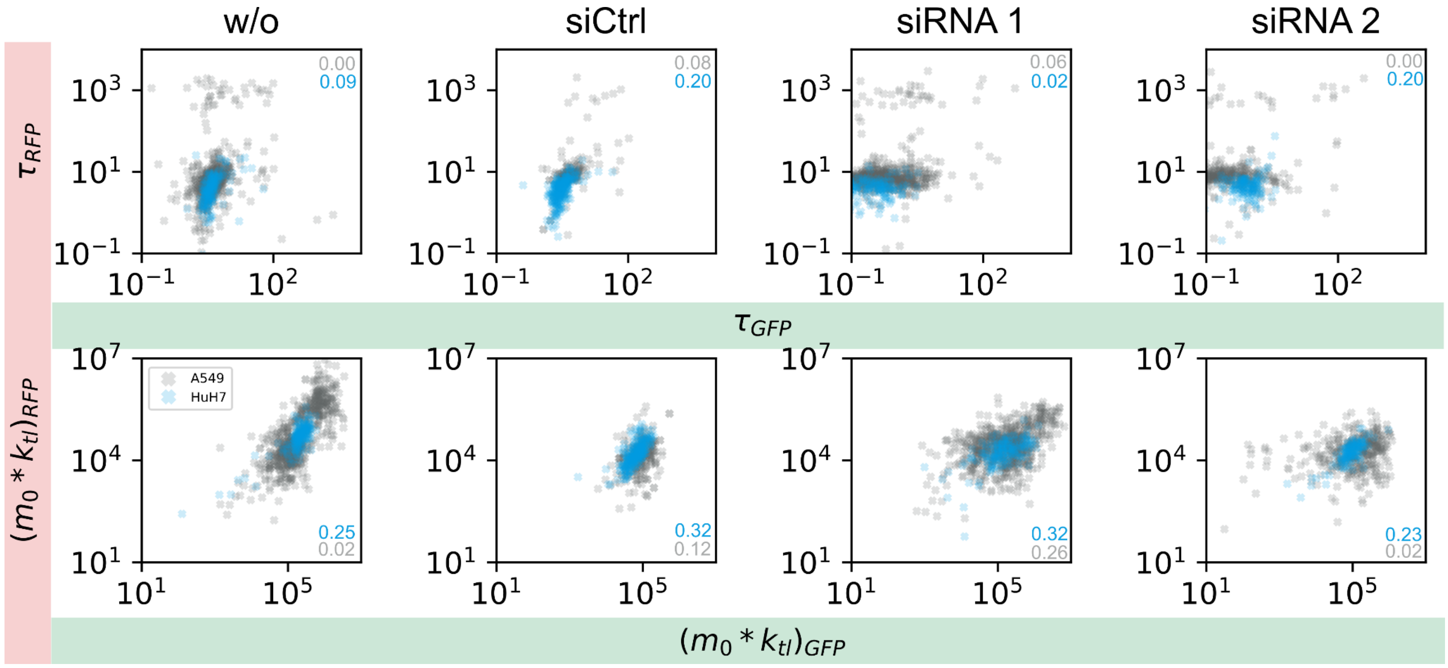


***Figure S5: eGFP-CayRFP correlation plots for stability τ and initial expression rate $m_0$*$k_{tl}$:*** *Top row: single-cell scatter plots for stability of eGFP and CayRFP mRNA. Bottom row: single-cell scatter plots of initial expression rates of eGFP and CayRFP mRNA. A weak correlation is expected because of a dominating effect of the initial mRNA amount $m_0$, which is equal for CayRFP and eGFP due to co-encapsulation. Gray crosses correspond to A549 cells, blue crosses to HuH7 cells. Values in the graphs show corresponding $R^2$ values.*

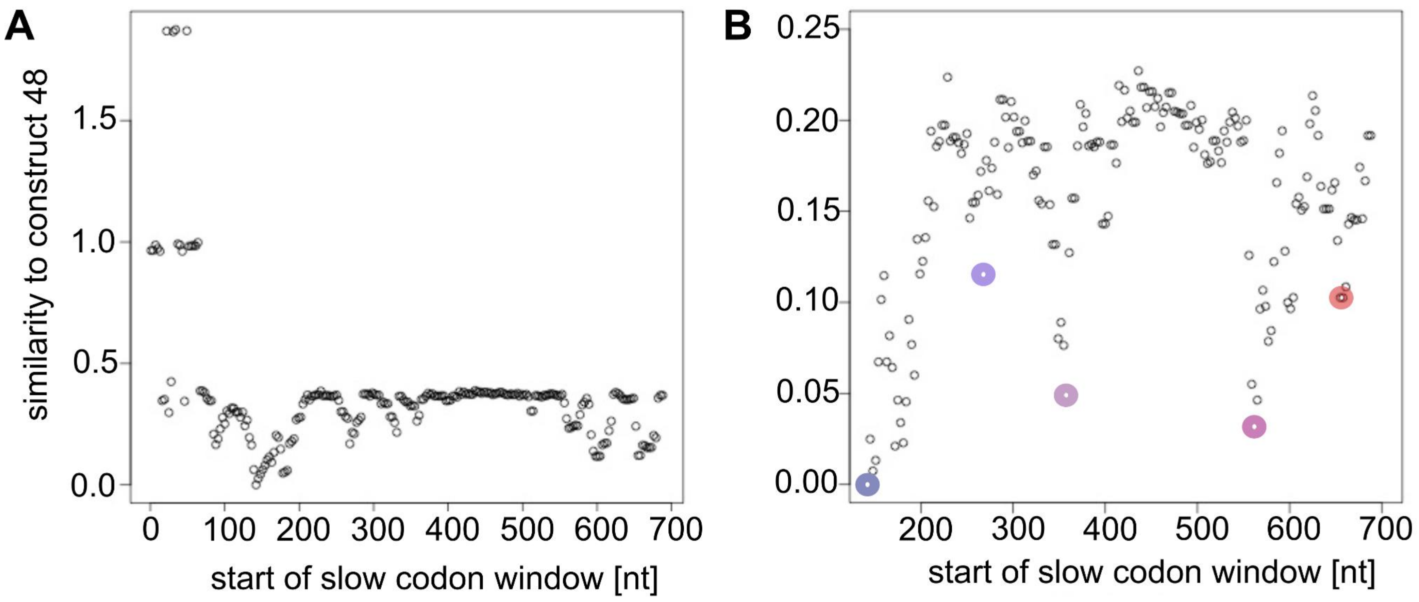


***Figure S6: Similarity to eGFP mRNA construct with slow-codon window starting from codon position 48:*** *We simulated230 versions of eGFP mRNA by moving a 10-codon window along the open reading frame and exchanging all codons in the window with their respective slowest (most non-optimal) synonymous alternatives based on codon-specific translation rates for human cells published in Trösemeier et al. [2]. Out of these 230 sequences, we first selected the construct with a slow-codon window directly downstream of the binding site of siRNA 1 (codon position 48 ff, "construct 48"). Next, we identified 4 additional constructs with slow-codon windows distributed homogeneously along the ORF and an overall rate of protein expression similar to that of construct 48 as computed by*

*simulations using OCTOPOS. (a) Similarity in protein expression rate to construct 48 for all 230 sequences. We observed high variations for sequences with slow-codon windows in the beginning of the ORF. (b) Zoom for better comparability. Selected constructs are marked with colored dots.*

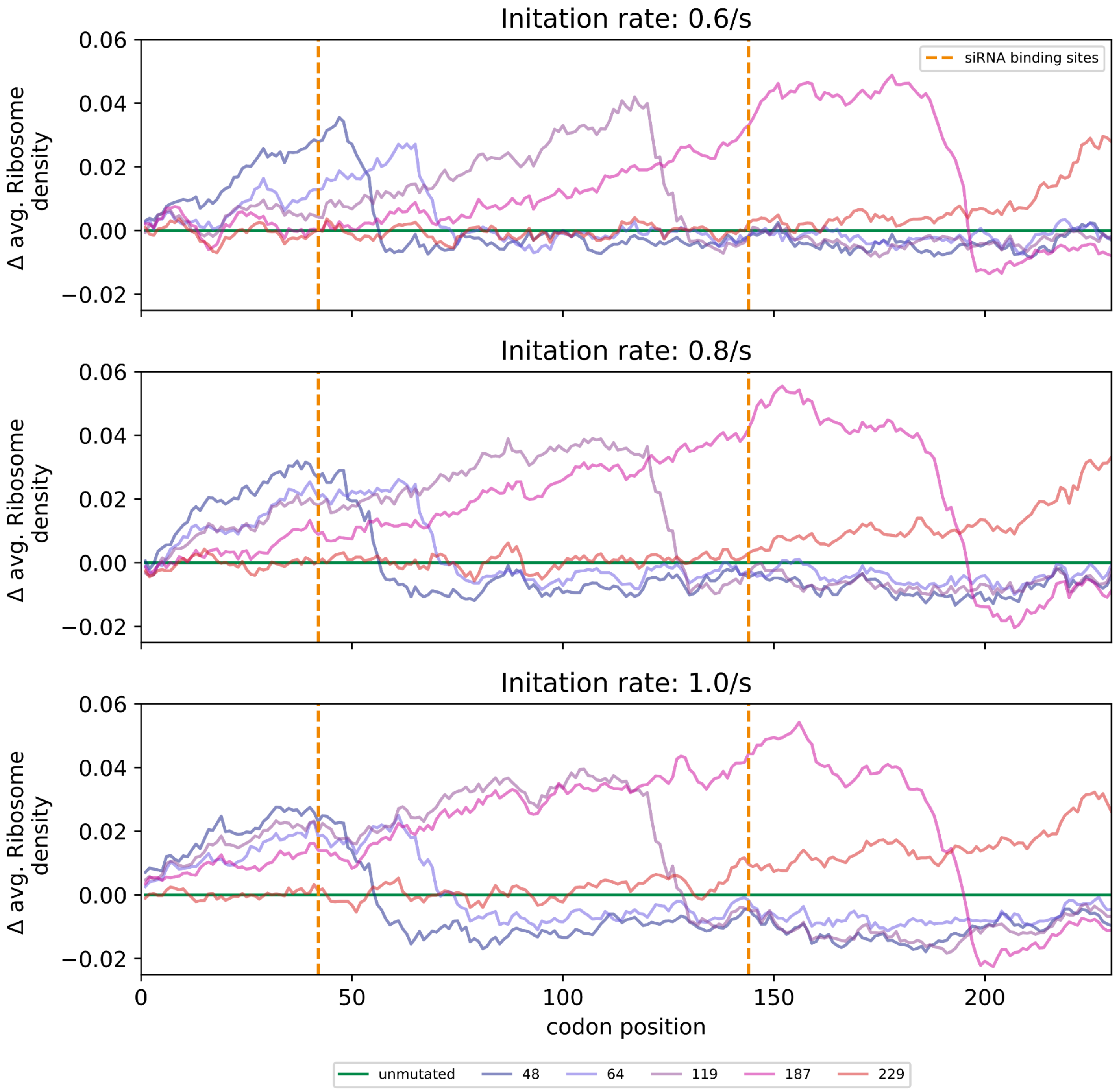


***Figure S7: Dependence of simulated ribosome density on initiation rate:*** *Initiation rates of 0,6, 0.8 and 1.0 per second are used as parameters for the translation simulation. Ribosome density was averaged over 10 codons for different eGFP mRNA constructs with slow-codon windows as depicted in* ***Fig. 3*** *in the main paper. Depicted is the difference in ribosome density compared to the unmutated eGFP mRNA.*

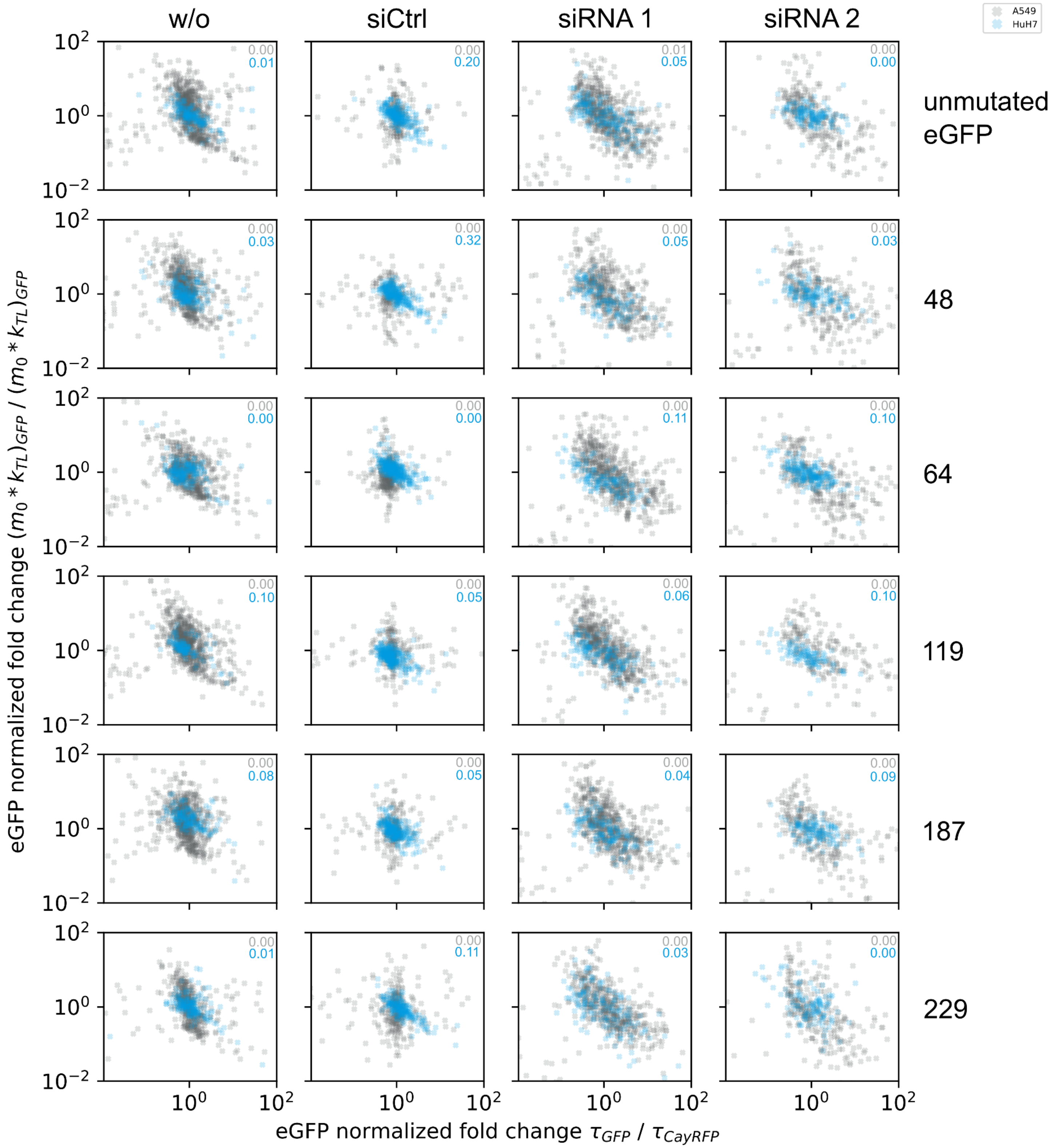


***Figure S8: Stability-expression scatter plots for all different mRNA constructs:*** *rows show data for the different eGFP variants and columns show experimental conditions: w/o: without siRNA addition, siCtrl: negative control siRNA, siRNA 1 and 2: addition of eGFP mRNA targeting siRNA. All eGFP data was normalized to the single-cell reference (GFP/CayRFP). Gray crosses correspond to A549 cells, blue crosses to HuH7 cells.* $R^2$ *values are shown in the plots in the respective color.*

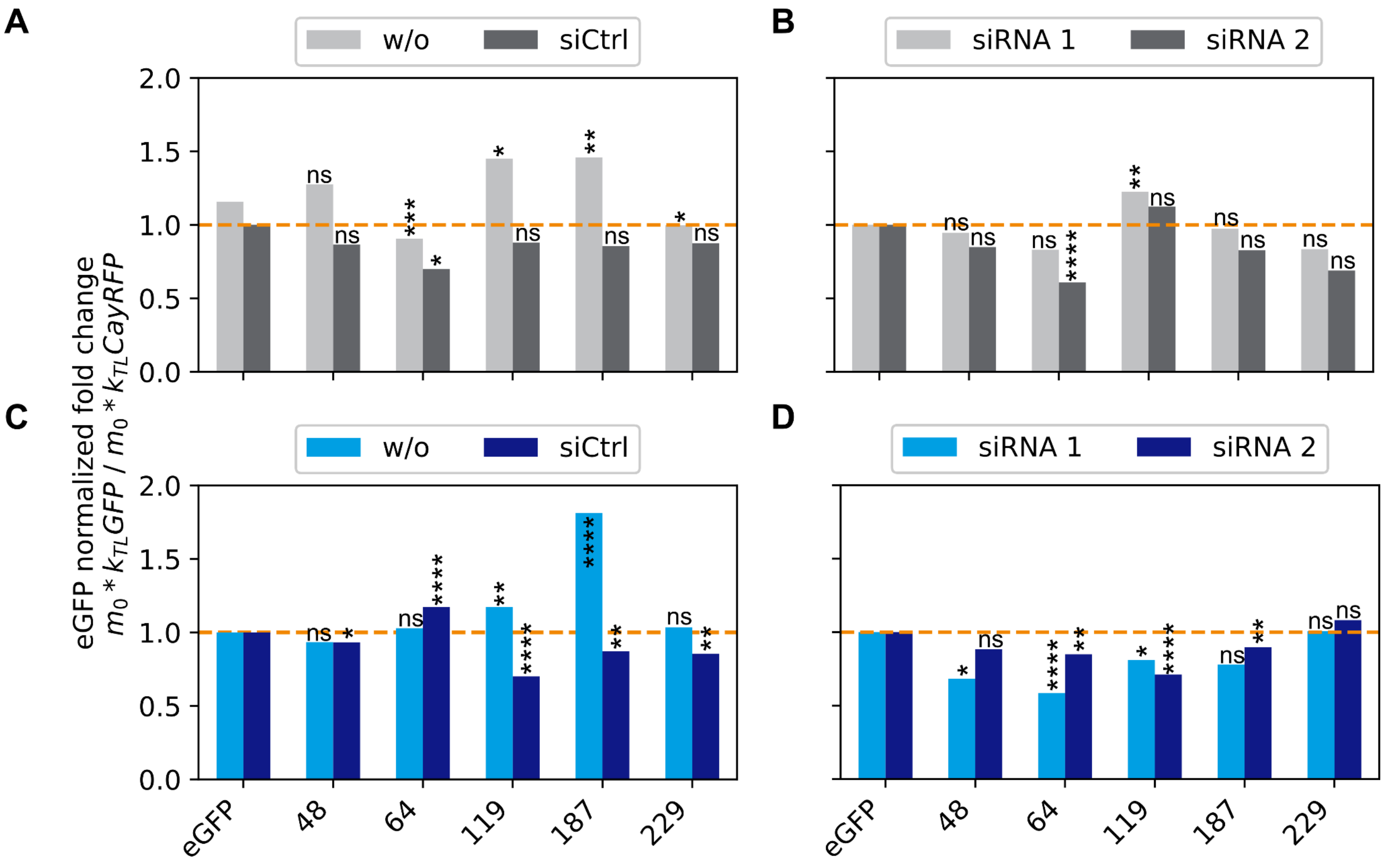


*Figure S9:* ***Effect of non-optimal codons on mRNA expression rate $k_{TL}$:*** *Expression rate of eGFP mRNA was determined by LISCA and normalized with equivalent values from reference CayRFP mRNA for each cell. (A)&(C) In absence of any siRNA, changes in expression rates were observed in both directions compared to unmutated eGFP in both A549 (gray) and HuH7 (blue) cells. (B) and (D) In the presence of negative control siRNA, again, changes in expression rates in both directions were observed in A549 cells, whereas HuH7 cells showed a trend towards decreased expression rates in presence of functional siRNA 1 and 2. Bars show median values of single cell data. Significance values by Mann-Whitney-U test.*

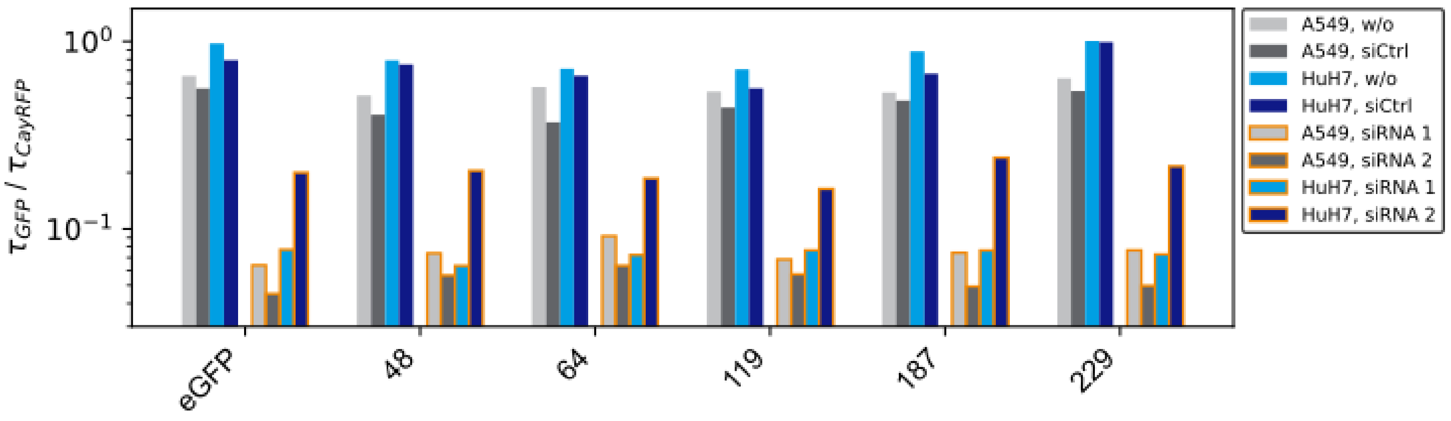


***Figure S10: Destabilization vs. stabilization – effect of slow-codon windows on mRNA stability:*** Stability of eGFP mRNA was determined by LISCA and normalized with equivalent values from reference CayRFP mRNA for each cell. Bars without borders show stability of tested mRNA constructs in absence of siRNA or with negative control siRNA. Bars with orange borders show stability of the same constructs but in presence of targeting siRNA. Stability in the presence of siRNA is reduced for all constructs but higher compared to eGFP mRNA. Gray bars correspond to A549 cells, blue bars to HuH7 cells. Normalized data and significance values can be found in **Fig. 4** of the main paper.

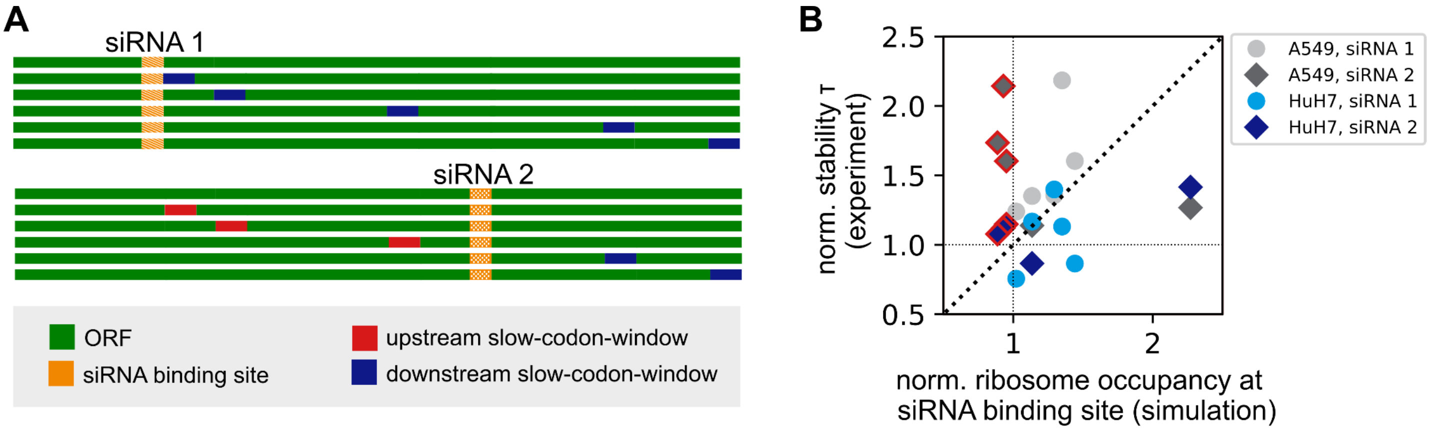


**Figure S11: Measured ribosome occupancy at siRNA binding site versus measured stability increase for the modified mRNA constructs.** *(A) Schematic representation of the eGFP construct and the 5 mutated constructs with inserted slow-codon windows. Red: inserted slow-codon-window upstream of the siRNA binding site, blue: downstream. siRNA 1 and 2 are indicated in yellow. (B) Measured fold change stability is plotted against the predicted ribosome density at the respective siRNA binding site. For slow-codon windows that are positioned downstream of the siRNA binding sites, the strength of stabilization correlates partially with the simulated ribosome densities at the siRNA binding sites. However, also mRNAs with non-optimal codon windows positioned upstream of the binding site of siRNA 2 (red bordered symbols) show increased relative stabilities. This indicated that the measured protective effect is not directly correlated with the predicted ribosome density at the siRNA binding site. All data is normalized to the stability of unmodified eGFP and the stability of the experiment in presence of siCtrl to account for the internal quality control decay.*

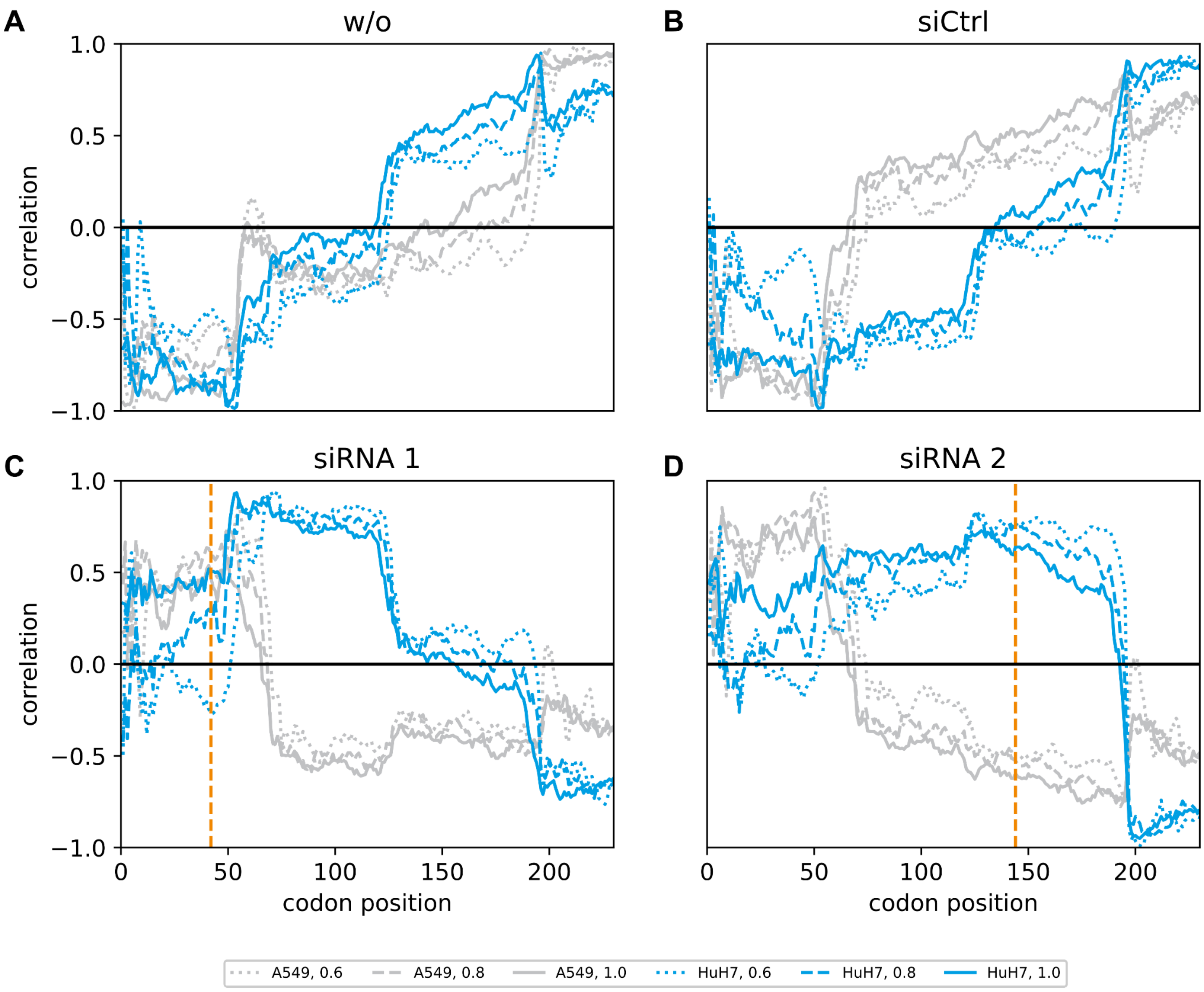


***Figure S12: Correlation coefficients of predicted ribosome density and measured fold changes of mRNA stability of modified eGFP constructs.*** *Different line-styles indicate different initiation rates (dotted: 0,6/s; dashed: 0.8/s; solid: 1.0/s), and orange dashed lines are positioned at the respective siRNA binding site. Gray curves correspond to A549 cells, blue to HuH7. (A) without siRNA, (B) with negative control siRNA, (C) with siRNA 1, (D) with siRNA 2.*

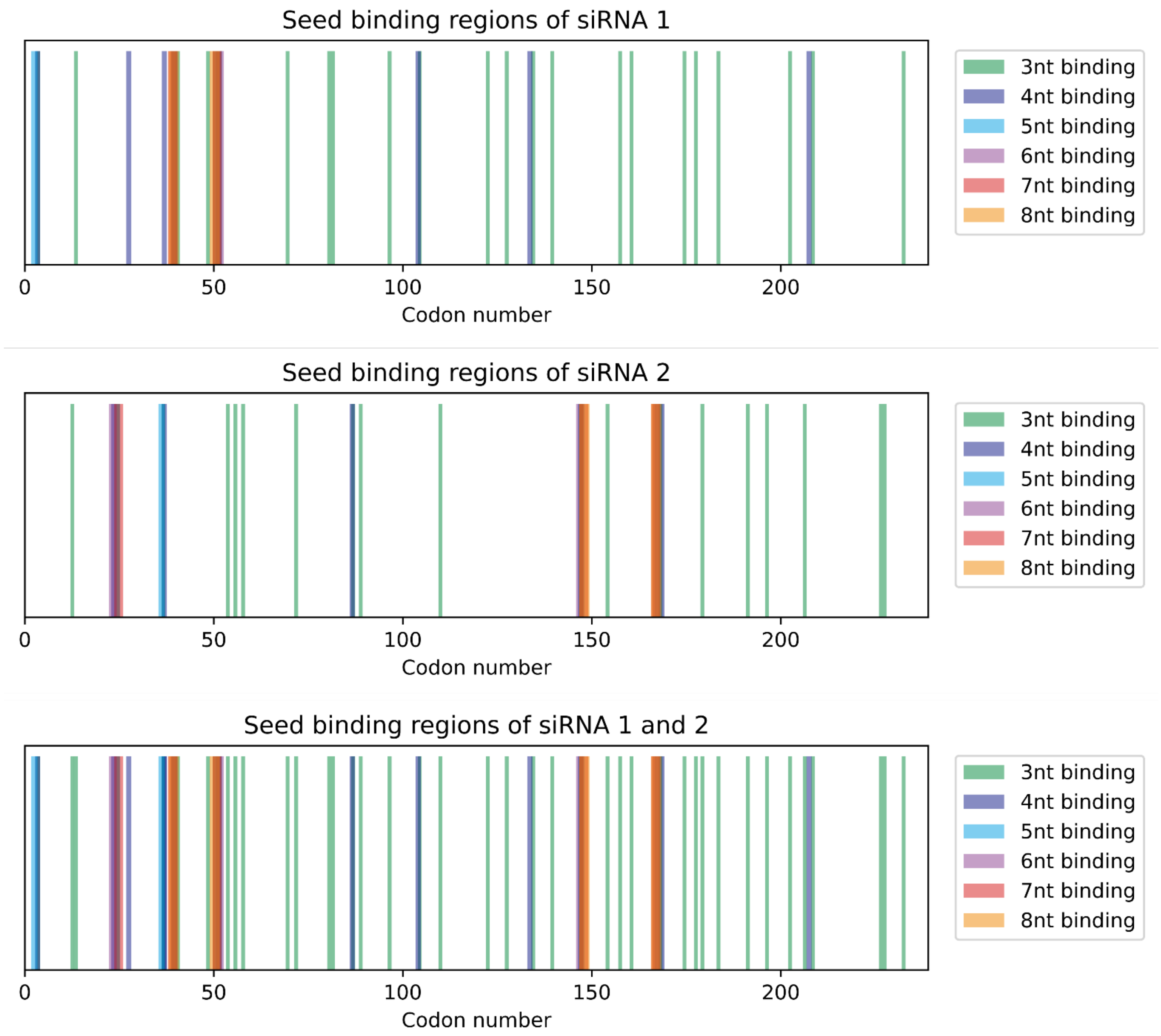


***Figure S13:*** *Regions on the ORF (valid for all mRNA sequences described in this work) to which the first 3 (green), 4 (blue), 5 (cyan), 6 (purple), 7 (red) or 8 nucleotides of the siRNA (seed region) are complementary.*

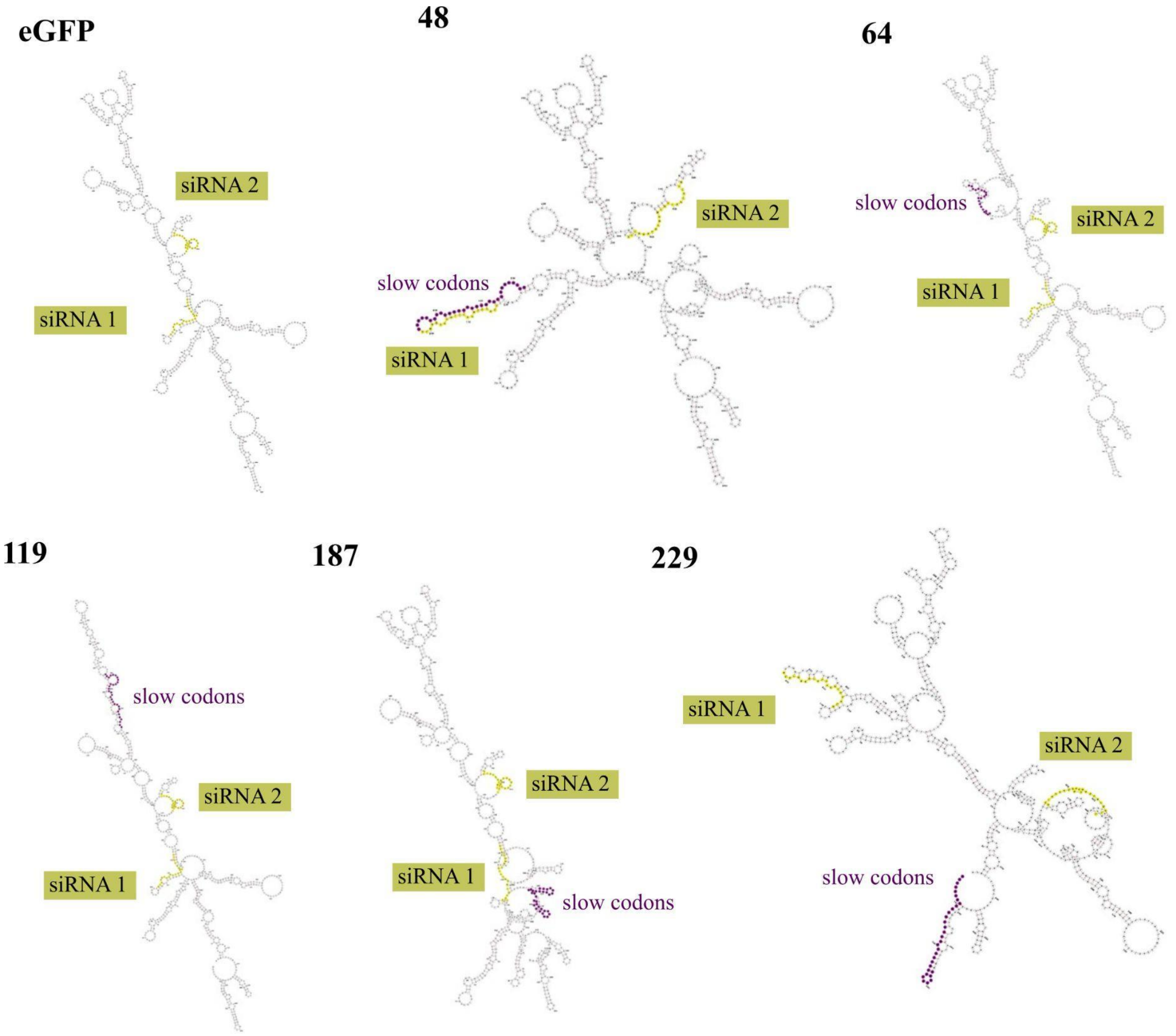


***Figure S14: Secondary structures of the mRNA constructs tested within this work:*** *yellow nucleotides mark the positions of both full siRNA binding sites; purple codons mark the synonymous slow-codon window. Numbers indicate codon-positions of the slow-codon windows. Structures were calculated with RNAfold via the minimal free energy option.*

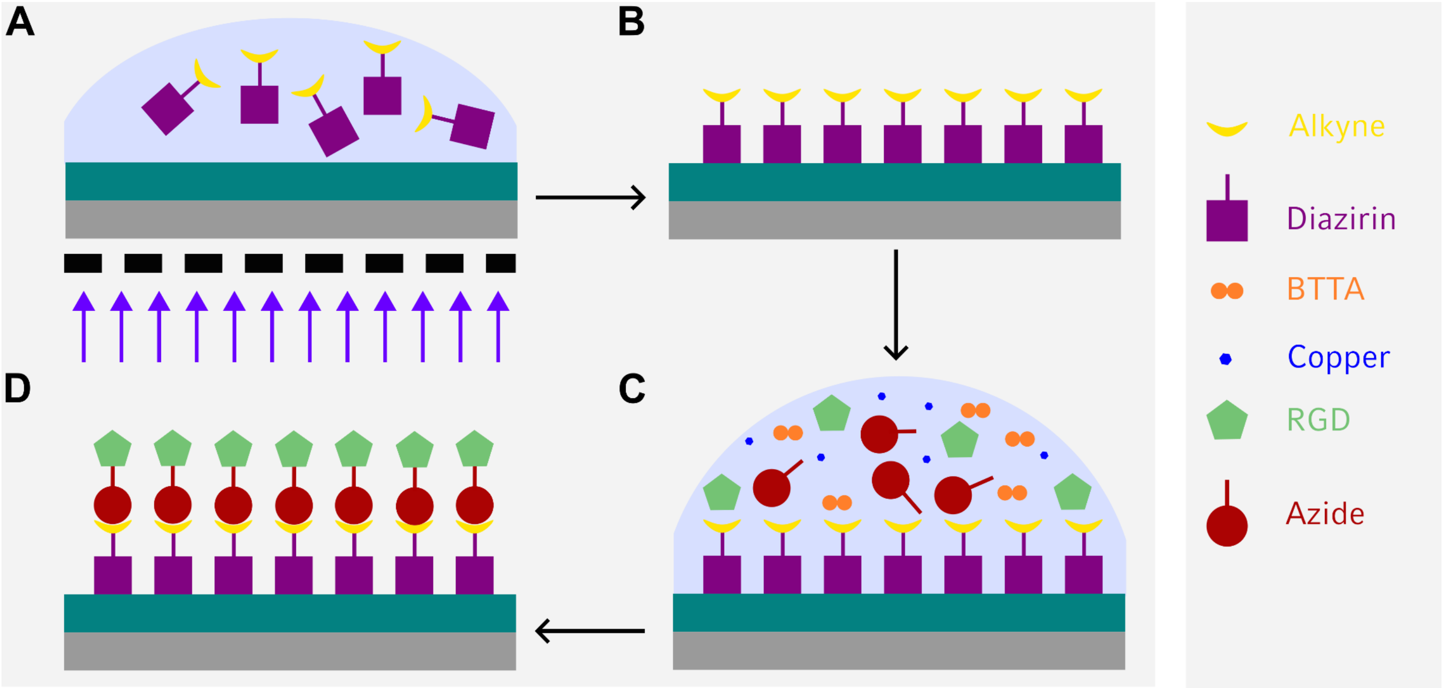


***Figure S15: Micropatterning for single cell assays:*** *(A) PVA coated ibidi µ-slide was coated with Diazirin-Alkyne and selectively illuminated with 365nm UV light. (B) Excess of Diazirin-Alkyne was washed away and (C) Cu-Click chemistry was employed to (D) selectively attach RGD.*

### mRNA and siRNA sequences

#### siRNA 1 (Dharmacon, GFP Duplex I siRNA):

Target sequence:  GCAAGCUGACCCUGAAGUUC

#### siRNA 2 (Dharmacon, custom production)

Target sequence: GCCACAACGUCUAUAUCAUUUU

#### eGFP sequence

ATGGTGAGCAAGGGCGAGGAGCTGTTCACCGGGGTGGTGCCCATCCTGGTCGAGCTGGACGGCGACGTAAACGGCCACAAGTTCA
GCGTGTCCGGCGAGGGCGAGGGCGATGCCACCTACGGCAAGCTGACCCTGAAGTTCATCTGCACCACCGGCAAGCTGCCCGTGCCC
TGGCCCACCCTCGTGACCACCCTGACCTACGGCGTGCAGTGCTTCAGCCGCTACCCCGACCACATGAAGCAGCACGACTTCTTCAAGT
CCGCCATGCCCGAAGGCTACGTCCAGGAGCGCACCATCTTCTTCAAGGACGACGGCAACTACAAGACCCGCGCCGAGGTGAAGTTC
GAGGGCGACACCCTGGTGAACCGCATCGAGCTGAAGGGCATCGACTTCAAGGAGGACGGCAACATCCTGGGGCACAAGCTGGAGT
ACAACTACAACAGCCACAACGTCTATATCATGGCCGACAAGCAGAAGAACGGCATCAAGGTGAACTTCAAGATCCGCCACAACATCG
AGGACGGCAGCGTGCAGCTCGCCGACCACTACCAGCAGAACACCCCCATCGGCGACGGCCCCGTGCTGCTGCCCGACAACCACTACC
TGAGCACCCAGTCCGCCCTGAGCAAAGACCCCAACGAGAAGCGCGATCACATGGTCCTGCTGGAGTTCGTGACCGCCGCCGGGATC
ACTCTCGGCATGGACGAGCTGTACAAG

#### Slow codon eGFP sequences

**48**

ATGGTGAGCAAGGGCGAGGAGCTGTTCACCGGGGTGGTGCCCATCCTGGTCGAGCTGGACGGCGACGTAAACGGCCACAAGTTCA
GCGTGTCCGGCGAGGGCGAGGGCGATGCCACCTACGGCAAGCTGACCCTGAAGTTCATATGTACAACAGGGAAGCTACCGGTACCC
TGGCCCACCCTCGTGACCACCCTGACCTACGGCGTGCAGTGCTTCAGCCGCTACCCCGACCACATGAAGCAGCACGACTTCTTCAAGT
CCGCCATGCCCGAAGGCTACGTCCAGGAGCGCACCATCTTCTTCAAGGACGACGGCAACTACAAGACCCGCGCCGAGGTGAAGTTC
GAGGGCGACACCCTGGTGAACCGCATCGAGCTGAAGGGCATCGACTTCAAGGAGGACGGCAACATCCTGGGGCACAAGCTGGAGT
ACAACTACAACAGCCACAACGTCTATATCATGGCCGACAAGCAGAAGAACGGCATCAAGGTGAACTTCAAGATCCGCCACAACATCG
AGGACGGCAGCGTGCAGCTCGCCGACCACTACCAGCAGAACACCCCCATCGGCGACGGCCCCGTGCTGCTGCCCGACAACCACTACC
TGAGCACCCAGTCCGCCCTGAGCAAAGACCCCAACGAGAAGCGCGATCACATGGTCCTGCTGGAGTTCGTGACCGCCGCCGGGATC
ACTCTCGGCATGGACGAGCTGTACAAG

**64**

ATGGTGAGCAAGGGCGAGGAGCTGTTCACCGGGGTGGTGCCCATCCTGGTCGAGCTGGACGGCGACGTAAACGGCCACAAGTTCA
GCGTGTCCGGCGAGGGCGAGGGCGATGCCACCTACGGCAAGCTGACCCTGAAGTTCATCTGCACCACCGGCAAGCTGCCCGTGCCC
TGGCCCACCCTCGTGACCACCCTAACATACGGGGTACAATGTTTCTCGCGGTACCCCGACCACATGAAGCAGCACGACTTCTTCAAGT
CCGCCATGCCCGAAGGCTACGTCCAGGAGCGCACCATCTTCTTCAAGGACGACGGCAACTACAAGACCCGCGCCGAGGTGAAGTTC
GAGGGCGACACCCTGGTGAACCGCATCGAGCTGAAGGGCATCGACTTCAAGGAGGACGGCAACATCCTGGGGCACAAGCTGGAGT
ACAACTACAACAGCCACAACGTCTATATCATGGCCGACAAGCAGAAGAACGGCATCAAGGTGAACTTCAAGATCCGCCACAACATCG
AGGACGGCAGCGTGCAGCTCGCCGACCACTACCAGCAGAACACCCCCATCGGCGACGGCCCCGTGCTGCTGCCCGACAACCACTACC
TGAGCACCCAGTCCGCCCTGAGCAAAGACCCCAACGAGAAGCGCGATCACATGGTCCTGCTGGAGTTCGTGACCGCCGCCGGGATC
ACTCTCGGCATGGACGAGCTGTACAAG

**119**

ATGGTGAGCAAGGGCGAGGAGCTGTTCACCGGGGTGGTGCCCATCCTGGTCGAGCTGGACGGCGACGTAAACGGCCACAAGTTCA
GCGTGTCCGGCGAGGGCGAGGGCGATGCCACCTACGGCAAGCTGACCCTGAAGTTCATCTGCACCACCGGCAAGCTGCCCGTGCCC
TGGCCCACCCTCGTGACCACCCTGACCTACGGCGTGCAGTGCTTCAGCCGCTACCCCGACCACATGAAGCAGCACGACTTCTTCAAGT

CCGCCATGCCCGAAGGCTACGTCCAGGAGCGCACCATCTTCTTCAAGGACGACGGCAACTACAAGACCCGCGCCGAGGTGAAGTTC
GAGGGCGACACCCTAGTAAACCGGATAGAGCTAAAGGGGATAGACTTCAAGGAGGACGGCAACATCCTGGGGCACAAGCTGGAGT
ACAACTACAACAGCCACAACGTCTATATCATGGCCGACAAGCAGAAGAACGGCATCAAGGTGAACTTCAAGATCCGCCACAACATCG
AGGACGGCAGCGTGCAGCTCGCCGACCACTACCAGCAGAACACCCCCATCGGCGACGGCCCCGTGCTGCTGCCCGACAACCACTACC
TGAGCACCCAGTCCGCCCTGAGCAAAGACCCCAACGAGAAGCGCGATCACATGGTCCTGCTGGAGTTCGTGACCGCCGCCGGGATC
ACTCTCGGCATGGACGAGCTGTACAAG

**187**

ATGGTGAGCAAGGGCGAGGAGCTGTTCACCGGGGTGGTGCCCATCCTGGTCGAGCTGGACGGCGACGTAAACGGCCACAAGTTCA
GCGTGTCCGGCGAGGGCGAGGGCGATGCCACCTACGGCAAGCTGACCCTGAAGTTCATCTGCACCACCGGCAAGCTGCCCGTGCCC
TGGCCCACCCTCGTGACCACCCTGACCTACGGCGTGCAGTGCTTCAGCCGCTACCCCGACCACATGAAGCAGCACGACTTCTTCAAGT
CCGCCATGCCCGAAGGCTACGTCCAGGAGCGCACCATCTTCTTCAAGGACGACGGCAACTACAAGACCCGCGCCGAGGTGAAGTTC
GAGGGCGACACCCTGGTGAACCGCATCGAGCTGAAGGGCATCGACTTCAAGGAGGACGGCAACATCCTGGGGCACAAGCTGGAGT
ACAACTACAACAGCCACAACGTCTATATCATGGCCGACAAGCAGAAGAACGGCATCAAGGTGAACTTCAAGATCCGCCACAACATCG
AGGACGGCAGCGTGCAGCTCGCCGACCACTACCAGCAGAACACCCCGATAGGGGATGGGCCGGTACTACTACCGGACAACCACTAC
CTGAGCACCCAGTCCGCCCTGAGCAAAGACCCCAACGAGAAGCGCGATCACATGGTCCTGCTGGAGTTCGTGACCGCCGCCGGGAT
CACTCTCGGCATGGACGAGCTGTACAAG

**229**

ATGGTGAGCAAGGGCGAGGAGCTGTTCACCGGGGTGGTGCCCATCCTGGTCGAGCTGGACGGCGACGTAAACGGCCACAAGTTCA
GCGTGTCCGGCGAGGGCGAGGGCGATGCCACCTACGGCAAGCTGACCCTGAAGTTCATCTGCACCACCGGCAAGCTGCCCGTGCCC
TGGCCCACCCTCGTGACCACCCTGACCTACGGCGTGCAGTGCTTCAGCCGCTACCCCGATCATATGAAGCAACATGATTTCTTCAAGT
CCGCCATGCCCGAAGGCTACGTCCAGGAGCGCACCATCTTCTTCAAGGACGACGGCAACTACAAGACCCGCGCCGAGGTGAAGTTC
GAGGGCGACACCCTGGTGAACCGCATCGAGCTGAAGGGCATCGACTTCAAGGAGGACGGCAACATCCTGGGGCACAAGCTGGAGT
ACAACTACAACAGCCACAACGTCTATATCATGGCCGACAAGCAGAAGAACGGCATCAAGGTGAACTTCAAGATCCGCCACAACATCG
AGGACGGCAGCGTGCAGCTCGCCGACCACTACCAGCAGAACACCCCCATCGGCGACGGCCCCGTGCTGCTGCCCGACAACCACTACC
TGAGCACCCAGTCCGCCCTGAGCAAAGACCCCAACGAGAAGCGCGATCACATGGTCCTGCTGGAGTTCGTGACCGCCGCCGGGATC
ACTCTCGGCATGGACGAGCTGTACAAG
